# ASD-FER: An Annotated Multi-Class Benchmark and Comparative Study for Facial Emotion Recognition in Children with Autism Spectrum Disorder

**DOI:** 10.64898/2026.10.14.754143

**Authors:** Shahana Yasmin Chowdhury, Bithi Banik, Tracey Ann Knaus, Christopher David Moore, Henry Fordjour Ansah, Huda Hammad, Pranish Ghimire, Md Tamjidul Hoque, Shreya Banerjee

**Affiliations:** Department of Computer Science LSU New Orleans, New Orleans, Louisiana, USA; Department of Psychology LSU New Orleans, New Orleans, Louisiana, USA; Kristiania University of Applied Sciences, Oslo, Norway

**Keywords:** Autism spectrum disorder, Facial emotion recognition, Children with autism, Transformer, Emotion Query Transformer (EQ-Former), Explainable AI, ASD-FER

## Abstract

Autism Spectrum Disorder (ASD) affects social interaction and communication, and automatically recognizing emotions from the facial expressions of children with ASD remains challenging. A key barrier to *facial emotion recognition* (FER) in children with ASD is the absence of ASD-specific datasets with proper emotion-level annotations. To address this, we present **ASD-FER**, a benchmark of 2,779 facial images of children with ASD annotated across six emotion categories by three independent annotators using majority voting. We evaluate nine models — CNN, SVM, AlexNet, ResNet34, ResNet50, MobileNetV2, and three backbone configurations of our proposed task-specific architecture, **EQFormer**: ViT, Swin-T, and a compact Hybrid CNN-Transformer (HybridT) — under stratified 10-fold cross-validation and a held-out 90/10 evaluation. ResNet50 led conventional models at 84.89% mean cross-validation accuracy; among transformer-based models, ViT led with 88.27%, followed by Swin-T at 88.02% and HybridT at 86.58%. On the held-out set, ViT achieved the highest overall accuracy (91.01%), while Swin-T obtained the best balanced accuracy (74.47%) and macro-F1 (0.7078); HybridT remained competitive at 89.57% accuracy with a fraction of the parameters. To support explainability, Grad-CAM visualizations highlighted attention on emotion-relevant facial regions — eyes, eyebrows, and mouth — with Swin-T the most spatially focused and the HybridT the most parameter-efficient (2.66M). Class imbalance and cross-dataset generalization remain open challenges.

## 1. Introduction

Autism Spectrum Disorder (ASD) is a complex neurodevelopmental disorder characterized by persistent differences in social communication, sensory processing, and behavioral flexibility (1). According to the World Health Organization, approximately 1 in 100 children worldwide is diagnosed with ASD, with prevalence estimates varying considerably across studies and regions (2). One of the most prominent manifestations of ASD is a marked difference in the production and recognition of facial expressions: individuals with ASD frequently exhibit atypical, ambiguous, or reduced expressions that diverge substantially from neurotypical patterns, making it harder for both human observers and automated systems to decode their emotional states (3; 4). Human emotion decoding itself relies heavily on the eye and mouth regions, with the relative importance of each shifting by emotion category (5; 6) — a finding that motivates the region-aware annotation and interpretability analysis adopted in this work. Early identification and monitoring of these expressive differences is critical; timely intervention during the formative years can substantially improve social communication outcomes (7).

The ability to automatically and reliably decode emotions from facial cues has broad clinical relevance in ASD care. Therapists, caregivers, and educators rely on emotional feedback to calibrate therapy sessions, assess engagement, and detect distress signals such as meltdown precursors. Manual observation is time-consuming, subjective, and infeasible at scale. Automated Facial Expression Recognition (FER) systems offer a promising complement to manual observation, enabling objective, continuous, and unobtrusive monitoring in naturalistic settings such as therapy rooms, classrooms, and homes (8).

Despite considerable advances in general-purpose FER driven by deep learning, applying these systems directly to ASD populations presents three interrelated challenges, as enumerated below, that are rarely addressed in combination:

1. **Atypical Emotional Presentation**: the facial Action Unit (AU) configurations produced by ASD individuals frequently differ from prototypical expressions used to train standard FER models, leading to systematic misclassification (3).
2. **Severe Class Imbalance**: ASD-specific datasets tend to contain substantially more instances of positive emotions (especially happiness) than negative or socially complex ones (fear, anger, surprise), skewing model learning and inflating macro accuracy while minority-class recall collapses (9).
3. **Limited Labeled Data**: collecting and annotating clinical-grade facial data from ASD children is resource-intensive and ethically constrained, resulting in datasets orders of magnitude smaller than those used to train general FER systems (10). These three challenges jointly limit the transferability of existing FER research to real-world ASD applications.

The rapid adoption of transformer architectures in computer vision such as Vision Transformer (11; 12), Data-efficient Image Transformers (DeiT) (13), and the Swin Transformer (14), has opened new possibilities for FER. Transformers capture long-range spatial dependencies and global context more effectively than convolutional neural networks (CNNs), which rely on local receptive fields (15). Several recent studies have explored transformer-based FER in general settings with strong results (9), yet their systematic evaluation in the specific context of ASD children’s emotion recognition remains scarce. Moreover, hybrid architectures that combine convolutional feature extraction with self-attention mechanisms have shown promise for compact, efficient deployment (16). Furthermore, transfer learning has emerged as the dominant paradigm for addressing limited labeled data in specialized domains. Pre-training on large-scale face or image datasets followed by fine-tuning on domain-specific corpora allows models to leverage rich prior representations while adapting to the distributional characteristics of the target population. For ASD facial emotion recognition specifically, pre-training on auxiliary ASD-related corpora (such as FERAC (17; 10)) before fine-tuning on the target dataset may better align learned representations with the atypical expression patterns observed in ASD children. However, the relative benefit of domain-specific pre-training versus large-scale general pre-training (*e.g.*, ImageNet-1K) has not been systematically benchmarked in this context.

A natural question is why another FER model is needed when existing systems already achieve over 90% accuracy on general FER benchmarks (9). The key challenge is that these results do not readily transfer to children with ASD, whose facial expressions are often more subtle and atypical. Existing ASD-specific FER studies are also limited in several important ways. Most rely on small datasets—for example, FERAC contains only 770 images across four emotion classes (17; 10)—which restrict model generalization. In addition, previous work has focused mainly on CNN-based methods, while the effectiveness of transformer architectures for ASD facial emotion recognition has received limited investigation. Even when transformers have been explored, they are rarely compared with CNNs under the same preprocessing pipeline, training strategy, and evaluation protocol, making it difficult to isolate the effect of the backbone architecture. To address this gap, this study presents a comprehensive evaluation of three complementary transformer paradigms–a lightweight HybridT, Swin-T, and ViT–under a unified experimental framework. We refer to this shared framework as **EQFormer** (Emotion Query Transformer); HybridT, Swin-T, and ViT denote its three backbone configurations. All three models share the same preprocessing, data augmentation, emotion-aware decoding, contrastive learning, and classification pipeline, differing only in their backbone feature extractors to enable a controlled architectural comparison. Furthermore, because overall accuracy alone can be misleading on highly imbalanced datasets, this work reports balanced accuracy and macro-F1 alongside overall accuracy to provide a more comprehensive evaluation, particularly for important minority emotions such as fear, sadness, and surprise. Finally, we introduce ASD-FER, an annotated benchmark dataset and evaluate nine machine learning and deep learning models—including classical machine learning, CNNs, and transformer-based architectures—under a unified training strategy and evaluation protocol. To improve model interpretability, we further employ Grad-CAM visualizations to examine whether the models’ predictions are grounded in emotion-relevant facial regions associated with emotion recognition.

## 2. Literature Review and Qualitative Comparison

The landscape of FER research relevant to ASD can be organized around three themes: (i) general-purpose FER benchmarks and architectures; (ii) ASD-specific binary classification (ASD vs. typically developing); and (iii) multiclass emotion recognition in ASD children. Table 1 provides a qualitative comparison of representative studies drawn from each category, summarizing their datasets, methods, tasks, best reported results, and key limitations. We discuss the most salient findings below.

**Table 1.** Qualitative comparison of representative facial expression and ASD-related visual recognition studies.

| Paper | Dataset | Subjects | Method | Task | Best Re- sult | ASD | Limitations |
| --- | --- | --- | --- | --- | --- | --- | --- |
| (18) | RAF-DB, AffectNet (used for FER) | General facial analysis | Swin-B + BDCA | unified 10-task | 88.24 (on FER) | No | Unequal representation across age, gender, and race across its datasets |
| (20) | Kaggle (2,940 images) | ASD vs TD | ResNet, VGG, MobileNet | Binary ASD Classification | 92% acc | Yes | Non-clinical dataset; no explainability. |
| (22) | East Asia ASD Children Facial | ASD vs TD | VGG16 | Binary ASD Classification | 95% acc | Yes | Single-race dataset; limited demographic diversity; |
| (10) | FERAC (770 images) | ASD children | Swin/ViT/DeiT | 4-class FER | 80.0% acc | Yes | Class imbalance; small dataset. |
| (3) | 4,595 images | ASD and TD | Random Forest + facial landmarks | Emotion production | 82.05 (TD); 66.43 (ASD) | Yes | Small ASD cohort; emotion confusion. |
| (24) | SSBD + Affect-Net | ASD videos | Perceiver | ASD + affect recognition | 75.2% (5-fold) | Yes | Low ASD accuracy; weak generalization. |
| (28) | ASD + TD; ASD images from Talaat (29) | ASD+TD | ViT (teacher) + Swin-T (student) with Uncertainty-Aware KD | 6-class, FER | 89.00% acc | Yes | Small dataset; limited demographic diversity;. |
| (27) | ASD children images from (30) | ASD | CNN, ResNet50, VGG16 | 6-class | 88%, 84%, 76% respectively | Yes | real-time images; augmentation before splitting raises potential leakage concern. |
| **Ours** | **2,779 images, 6 classes** | **ASD children** | **CNN, SVM, ResNet, AlexNet, MobileNet, HybridT, Swin-T, ViT** | **Multi-class FER** | **91.01%** | **Yes** | **Class imbalance; single dataset; cross-dataset validation needed** |

**General-purpose FER.** Systems trained on large datasets such as FER-2013, AffectNet, RAF-DB, and CK+ have achieved impressive benchmark performance. The comprehensive survey by Li and Deng (9) catalogs methods ranging from hand-crafted HOG and geometric features to CNN, GAN, GCN, and transformer pipelines, with leading models exceeding 90% accuracy on standard benchmarks. FaceXFormer (18) represents the state of the art in unified facial analysis, combining a Swin-B encoder with a bidirectional cross-attention decoder to achieve 88.24% accuracy on RAF-DB (19). However, none of these systems account for the atypical AU configurations characteristic of ASD, and none are validated on ASD populations—direct application therefore risks systematic bias introduced by exclusively neurotypical training data.

**ASD binary classification.** Several works utilize the publicly available Kaggle ASD dataset for binary ASD vs. typically developing (TD) classification (20; 21; 22). ASD-CNN (20) achieves 92% accuracy using ResNet50 but lacks demographic stratification and explainability. DL_BinClass (21) obtains comparable results with MobileNetV2 (92% accuracy, AUC 0.92). Critically, the ethnoracial bias study (22) demonstrates that classifiers trained on Caucasian cohorts degrade substantially when applied to East Asian populations, highlighting a fairness concern for clinical deployment. Beyond appearance-based approaches, Xia and colleagues (23) propose dynamic viewing pattern analysis using an Autoencoder + LSTM pipeline on 333 children, achieving 96.73% accuracy for ASD screening (the highest reported in this category), though this comes at the cost of requiring specialized eye-tracking hardware, limiting scalability in remote or low-resource settings. All of these works, however, frame the problem as binary and do not model the space of individual emotions, limiting their utility for fine-grained emotional monitoring.

**Multi-class emotion recognition in ASD children.** This sub-area is considerably less explored. The FERAC dataset (10), containing 770 images across four emotion classes from autistic children, represents one of the few ASD-specific FER benchmarks. The authors of (10) find that Swin achieves the best performance (80.0% accuracy, F1 0.789) but suffers from extreme class imbalance—the fear class contains only 49 instances compared to 463 for joy. Grossard et al. (3) further demonstrate that ASD children produce more ambiguous and less socially meaningful facial expressions than TD peers, with Random Forest classifiers achieving 82.05% accuracy on TD children but showing significant emotion confusion in the ASD cohort — underscoring the inadequacy of neurotypical-trained FER models for this population. Extending this to video, Dia and colleagues (24) apply a CNN + Perceiver architecture on SSBD and AffectNet for continuous affect recognition in ASD children, but achieve only 74.5% accuracy with limited generalization across subjects. The CALMED dataset (25) provides multimodal video and audio annotations for four ASD children using a zone-based emotion model, but no classification model has been published on it. The real-time FER system of (26) targets mobile deployment to assist ASD individuals in recognizing others’ expressions, yet it is trained entirely on FER-2013 with minimal ASD-specific validation. More recently, Hamawy *et al.* (27) apply CNN, ResNet50 and VGG16 to a six-class ASD emotion dataset, reporting 90.0%, 88.0% and 78.0% accuracy respectively, though their evaluation relies on augmented test data. Christina and Sweetlin (28) propose a contextual multi-head transformer with uncertainty-aware distillation for ASD emotion recognition, reporting 89.0% accuracy, but their work stops short of releasing a benchmark dataset or conducting a systematic comparison between CNNs and transformers **Explainability in FER.** Explainability has received growing attention in general-population FER. Grad-CAM, originally introduced by Selvaraju *et al.* (31) as a gradient-based technique for visualizing the image regions driving a CNN’s prediction, has since become a standard tool for auditing whether FER models attend to expression-relevant facial regions rather than background or spurious cues. Applying this technique across FER-2013, RAF-DB, and AffectNet, Gebele *et al.* (32) combine Grad-CAM with Guided Backpropagation and find that the features learned by CNN-based FER models do not always align with expert emotion-psychology theory, underscoring that high accuracy alone does not guarantee that a model has learned clinically or psychologically meaningful cues. However, such analyses are confined to typically-developing (TD) populations and standard benchmarks. None of the ASD-specific studies summarized in Table 1 report any form of spatial explainability, leaving open whether existing ASD FER classifiers are exploiting facial regions with clinical relevance or are instead fitting to dataset artifacts and source-image biases–a concern that is particularly acute given the internet-sourced, diagnostically-unverified nature of most ASD facial datasets discussed above.

Taken together, these strands of work reveal that general-purpose FER systems are not validated on ASD populations, ASD binary-classification studies do not model individual emotions, and the handful of multi-class ASD FER studies are constrained by small, imbalanced datasets and narrow architectural comparisons.

### 2.1. Research Gaps

These observations translate into five principal gaps that motivate the present work:

1. **Absence of systematic transformer benchmarking for ASD FER.** While transformers have demonstrated superiority over CNNs on general FER benchmarks, no study has rigorously compared full transformer architectures (e.g., Swin-T) against classical ML and CNN baselines specifically for multi-class emotion recognition in ASD children under a single, matched preprocessing/training pipeline with stratified cross-validation and an independent held-out evaluation. Radočaj and Martinović (10) compare several transformer variants but do not include CNN baselines, domain-specific pre-training, or a held-out generalization test.
2. **Lack of domain-specific pre-training analysis.** The relative contribution of pre-training on ASD-relevant corpora (FERAC + FER-Autism (17; 33)) versus large-scale general pre-training (ImageNet-1K) has not been disentangled, leaving practitioners without evidence-based guidance on transfer learning strategies for this domain.
3. **Insufficient explainability.** With few exceptions, existing ASD FER work does not provide spatial explanations (*e.g.*, Grad-CAM) to verify that models attend to meaningful facial regions, limiting interpretability and clinical trustworthiness.
4. **Minority emotion under-recognition.** Fear, surprise, and sadness—important signals in ASD monitoring—are systematically underperformed in existing models due to severe class imbalance, yet targeted balancing strategies tailored to ASD datasets remain underexplored.
5. **Absence of a multi-class, emotion-annotated ASD facial dataset at scale.** Existing ASD-specific resources are either binary-labeled (ASD vs. TD) or, where emotion-annotated, extremely small (e.g., FERAC’s 770 images across four classes), limiting the statistical power of any downstream benchmarking effort.

### 2.2. Contributions

In response to the identified gaps, this paper makes the following contributions:

1. **Multi-Class benchmark.** Addressing the dataset-scale gap identified above, we extract initially 2,784 ASD-only facial images from an existing ASD screening dataset (later reduced to 2,779 after excluding five non-consensus annotations, as detailed in Section 3), setting aside the typically developing (TD) samples, to build a dedicated benchmark for studying emotion expression in children with ASD. We also annotate each image with one of six fine-grained emotion labels, moving beyond the source dataset’s binary diagnostic label (ASD/TD) and enabling detailed FER analysis.
2. **Unified comparative benchmark.** We benchmark nine models—a baseline CNN, SVM, ResNet34, ResNet50, AlexNet, MobileNetV2, a compact HybridT, Swin-T and ViT — under a consistently applied evaluation protocol consisting of stratified 10-fold cross-validation and an independent 90/10 held-out test set for final performance estimation. To our knowledge, this is the first study to directly compare traditional, CNN-based, and transformer architectures for *multi-class* ASD emotion recognition under such a controlled protocol. We name the shared query-based decoding pipeline underlying HybridT, Swin-T, and ViT as **EQFormer**, and show that its architectural benefits hold consistently across all three backbone configurations on the ASD-FER dataset.
3. **Domain-specific pre-training strategy.** We pre-train the CNN-based models (initialized from ImageNet-1K) and, separately, the from-scratch HybridT, on a combined FERAC + FER-Autism corpus before fine-tuning on ASD-FER, while Swin-T and ViT use their respective ImageNet-pretrained backbones without further domain-specific pretraining. This setup allows us to examine the performance of ASD-relevant pre-training within the proposed benchmark, while recognizing that the pre-training configurations are not strictly identical across all architectures.
4. **Efficiency–accuracy trade-off analysis.** By including a HybridT model with only 2.66M parameters, alongside Swin-T (44.7M parameters) and ViT (102.95M parameters), we provide evidence-based guidance for practitioners facing deployment constraints. The HybridT model achieves 89.57% accuracy and a macro-F1 score of 0.6924, remaining competitive with Swin-T, which attains 90.29% accuracy and a macro-F1 score of 0.7078, and ViT, which achieves 91.01% accuracy and a macro-F1 score of 0.7024, while requiring only a fraction of the parameter count.
5. **Grad-CAM explainability.** We apply Grad-CAM to all transformer models to verify that predictions are anchored in meaningful facial regions (eyes, eyebrows, mouth), providing qualitative interpretability evidence that can support inspection of whether predictions are associated with meaningful facial regions.
6. **Minority-class performance analysis.** We provide detailed per-class F1 scores and balanced accuracy metrics that expose the persistent difficulty of recognizing low-prevalence emotions (Fear, Surprise, Sadness) and frame these as concrete targets for future work.

## 3. The ASD-FER Dataset

### 3.1. Source Data and Motivation

The source data for ASD-FER originates from a publicly available Kaggle dataset titled Facial_data_ASD (34), uploaded in March 2022. The dataset was originally constructed as a binary-class resource containing facial images of children with ASD and typically developing (TD) children, collected through internet searches across autism-related websites and social media communities. It has been used in prior work exclusively for binary ASD vs. TD classification tasks — that is, to distinguish children with ASD from neurotypical peers based on facial appearance alone. To the best of our knowledge, no prior study has transformed this dataset into a multi-class facial emotion recognition benchmark; the emotion states of the children depicted have never been annotated. The facial images displayed in Figures 3, 4, 5, 12, 13, and 14 are derived from the same Kaggle Facial_data_ASD (34).

For the present work, we extracted the 2,784 ASD-only facial images from this dataset, setting aside the TD images entirely. Our objective was not to perform ASD diagnosis — a task the source dataset was designed for — but rather to construct the first annotated benchmark for studying how children with ASD express facial emotions across multiple categories. This distinction is critical: while the source dataset carries only a binary diagnostic label (ASD or TD), our work introduces six fine-grained emotion class labels for each image.

The six emotion classes chosen for this study—*angry*, *fear*, *happy*, *neutral*, *sad*, and *surprise*-follow Ekman and Friesen’s widely adopted framework of basic emotions (35). This taxonomy is the most widely adopted scheme in FER research (9) and aligns with the emotion categories used in existing ASD-specific FER studies such as FERAC (10) and the Autistic Children Emotions dataset (29), enabling meaningful cross-study comparisons. Disgust was excluded because it is relatively rare, difficult to annotate reliably in children, and often confused with anger in low-resolution facial images (36).

**Ethical and data quality considerations:** Like most publicly available ASD facial datasets, the source images were gathered via internet search rather than clinical recruitment, so diagnoses are unverified and parental consent is undocumented — limitations that led to the retraction of a comparable Kaggle-based study in *PLOS ONE* in December 2025 (37). We use this dataset transparently, as no emotion-annotated ASD facial resource currently exists. ASD-FER is a first step toward filling that gap, and we encourage future work to build on it with ethically sourced, clinically verified data.

### 3.2. Annotation Methodology

#### 3.2.1. Annotator Selection

Three raters involved in the study labeled each image. The raters included a psychology associate professor and two computer science graduate students. One of the CS graduate students is also a parent of a child with ASD. Each rater independently viewed each image and noted the emotion (angry, fear, happy, neutral, sad, or surprise) displayed by the child. While the entire face as a whole was used to determine the emotion, the eyes and eyebrows were key areas in making a decision, followed by the mouth region. This is consistent with human psychophysical findings that observers rely heavily on the eye and mouth regions when decoding facial expressions (5; 6). On images with any discrepancy in the label between the three raters, consultation between the three raters took place in order to come to a consensus. There were five images where raters were not able to determine the main emotion displayed or a consensus could not be reached and these images were therefore excluded from the study. Fig. 1 shows the annotation pipeline for the ASD-FER dataset.

**Figure 1:**
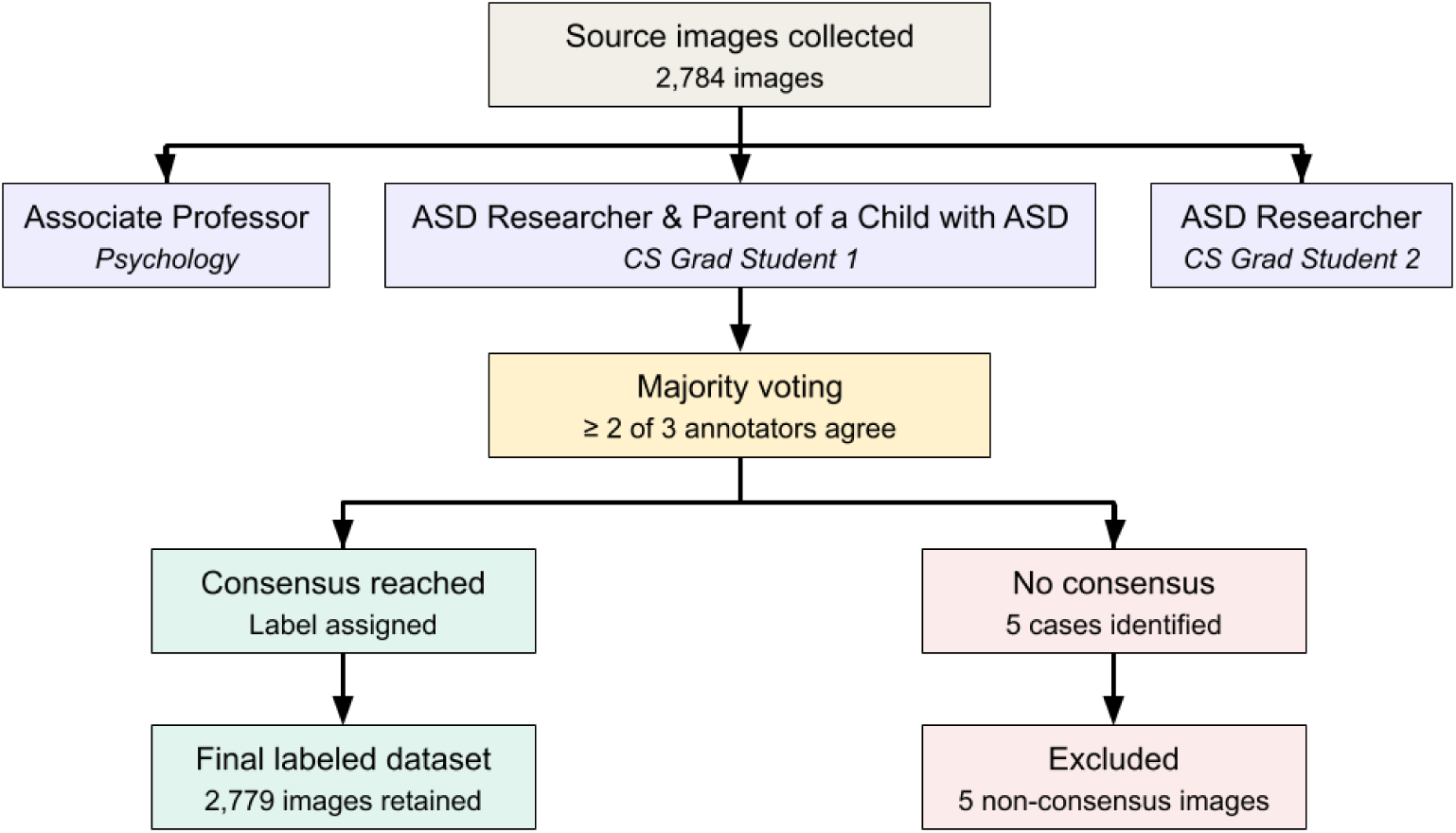
Annotation pipeline for the ASD-FER dataset. Each source image was independently labeled by three annotators (a psychology associate professor and two computer science graduate students) and assigned a final label via majority voting (≥ 2 of 3 agreement). The 5 non-consensus images were excluded, yielding 2,779 labeled images.

#### 3.2.2. Annotation Process

The labeling procedure was done by accessing the research group Microsoft OneDrive image folders. Emotion labels were then added using the comments section of each image. After all images were labeled by all three raters, these labels were then added to a spreadsheet for final label assessment. The raters had a choice of six emotion labels.

#### 3.2.3. Consensus and Final Label Assignment

Following the consultation process described in Section 3.2.1, final labels were assigned using a majority voting rule: an image’s label was accepted once at least 2 of the 3 annotators agreed on the same emotion class, whether that agreement was reached on initial labeling or after discussion. Images for which all three annotators could not reach agreement on a single emotion class even after consultation were treated as non-consensus and excluded from the dataset.

This process identified 5 non-consensus images out of the original 2,784 collected images, which were removed, yielding the final retained dataset of 2,779 original images used for all subsequent stratified 10-fold cross-validation and augmentation described in Section 3.4.2.

#### 3.2.4. Inter-Rater Agreement

To assess label reliability, we computed inter-rater agreement on the 2,687 images (96.5%) where all three annotators selected one of the six predefined emotion categories, based on each annotator’s initial label prior to consultation. Unanimous agreement across all three raters on this initial pass was 78.9%. Pairwise Cohen’s *κ* (38) ranged from 0.65 to 0.83 (mean *κ* = 0.73), and Fleiss’ *κ* (39) across all three raters was 0.73, indicating *substantial agreement* under the Landis and Koch benchmark (40). Images without initial unanimous agreement, including the remaining 97 images (3.5%) where at least one annotator gave a free-text or ambiguous response, went through the consultation process described in Section 3.2.1 and were resolved via the majority-voting rule in Section 3.2.3. After consultation, only 5 images had no resolvable consensus and were excluded from the final dataset, yielding the final 2,779 images used in ASD-FER.

### 3.3. Dataset Statistics and Distribution

The ASD-FER dataset contains 2,779 facial images across six emotion classes: Angry, Fear, Happy, Neutral, Sad, and Surprise (34). As shown in Table 2, the dataset is highly imbalanced, with Happy and Neutral accounting for nearly 89% of all images, while the remaining four emotions comprise only about 11%. This imbalance makes emotion recognition particularly challenging, especially for the minority classes.

**Table 2.** Per-class image count and percentage distribution in ASD-FER.

| Emotion | Image Count | Percentage (%) |
| --- | --- | --- |
| Angry | 112 | 4.03 |
| Fear | 82 | 2.95 |
| Happy | 1,717 | 61.79 |
| Neutral | 758 | 27.28 |
| Sad | 51 | 1.84 |
| Surprise | 59 | 2.12 |
| **Total** | **2,779** | **100.00** |

To provide a fair assessment of model performance, balanced accuracy and macro-F1 were used alongside overall accuracy, as these metrics better reflect performance across both majority and minority classes.

Fig. 2 shows the class distribution and confirms that the stratified 10-fold cross-validation protocol (41; 42) preserves the original class proportions in every fold, ensuring consistent representation of all emotion classes during training and evaluation.

**Figure 2:**
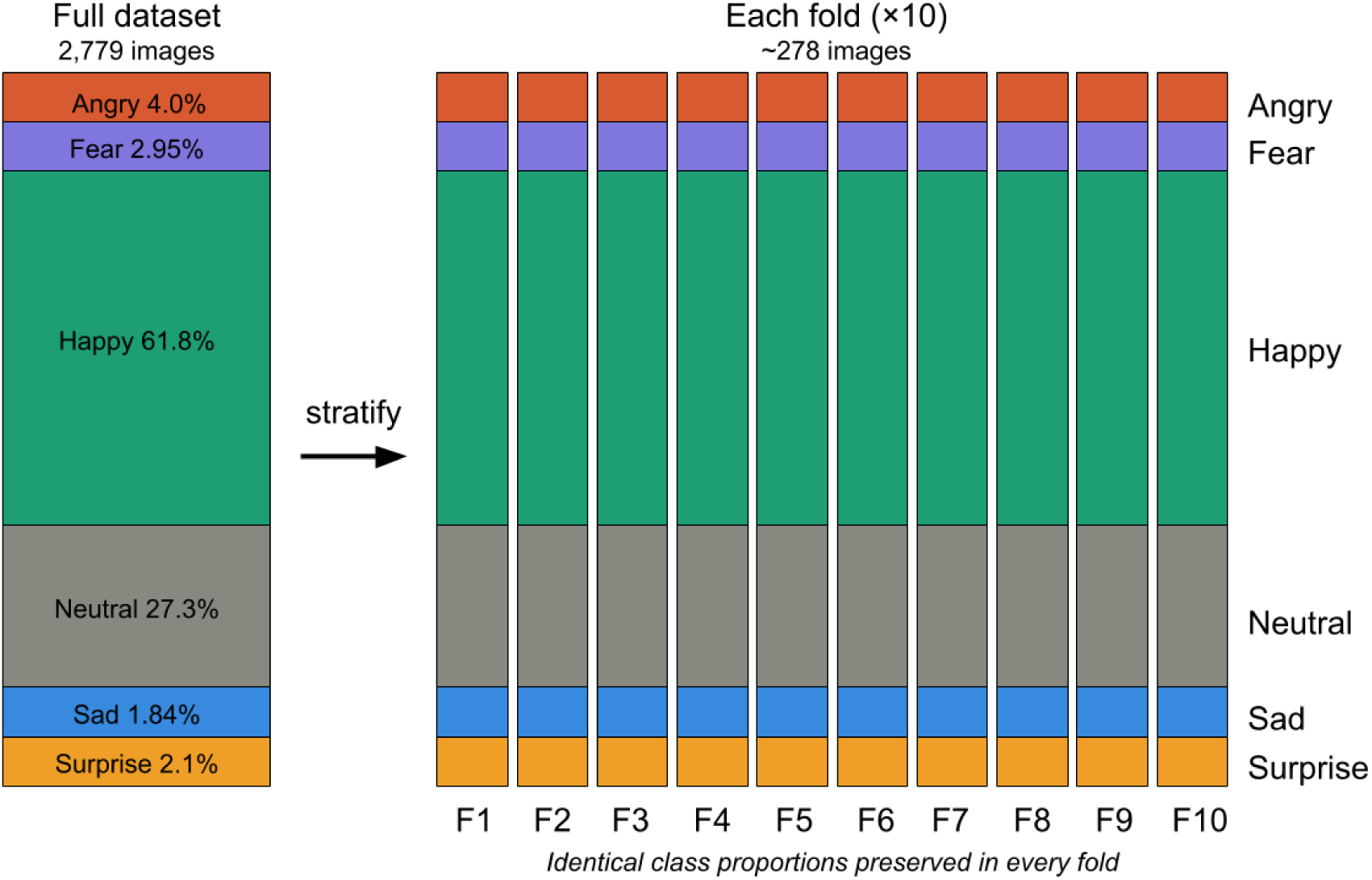
The stacked bars confirm that stratified 10-fold cross-validation preserves the severe class imbalance — dominated by Happy (61.8%) and Neutral (27.3%), with minority classes collectively below 11% — identically across all 10 folds, ensuring consistent representation of rare emotion classes in every train/validation/test split.

**Figure 3:**
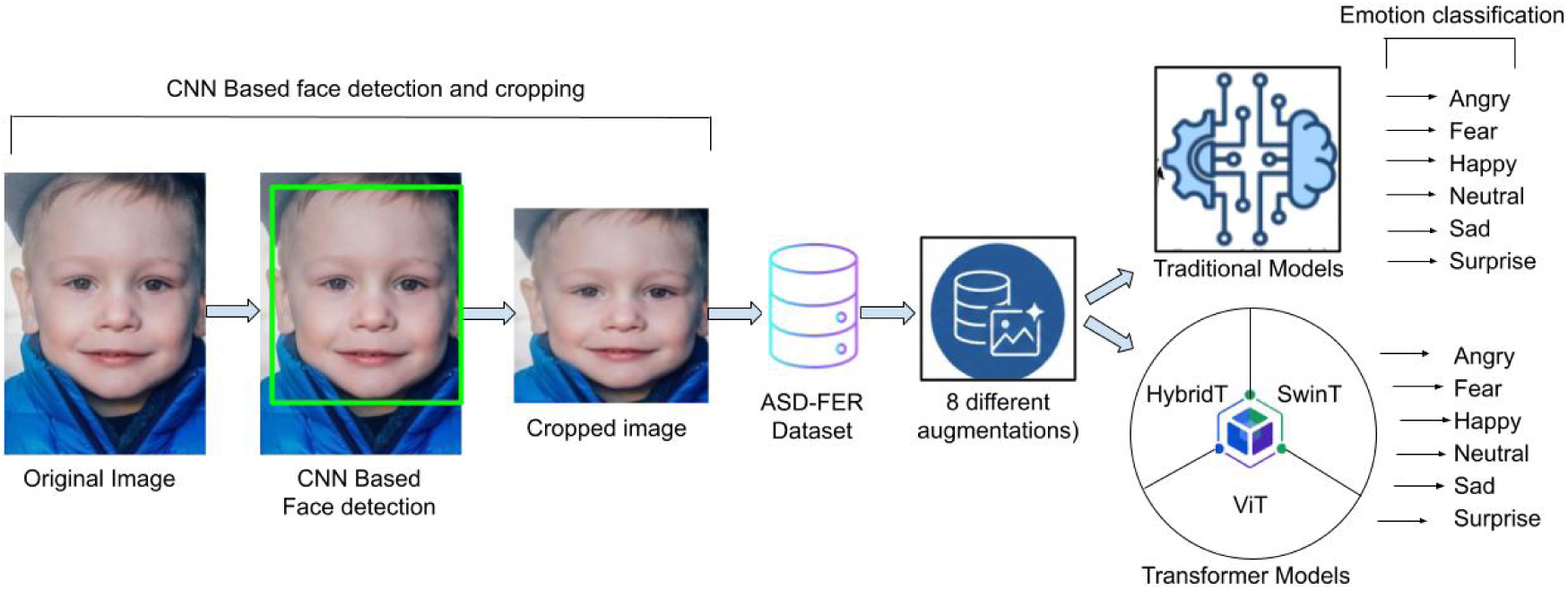
Preprocessing and augmentation workflow for emotion classification on the ASD-FER dataset. The facial images are derived from the publicly available Kaggle dataset, Facial_data_ASD, by Md. Shafiul Alam (34)

### 3.4. Data Preprocessing and Augmentation

#### 3.4.1. Data Preprocessing

Data Preprocessing is an important technique because it prepares the raw data for analysis and modeling (43). It also helps to extract relevant features, eliminate unnecessary information, and enhance model accuracy. As a part of data preprocessing, several techniques were applied to standardize and enhance the Kaggle dataset. As a first step, we used an open-source library for CNN-based face detection for images (44), which detects and crops the face from the image to isolate the facial features (see Fig. 3). In the next step, all cropped images were resized to 224 × 224 pixels to standardize the input size. We used preprocessed 2,779 images for our study.

To prevent augmentation-derived leakage across data partitions, dataset splitting was performed at the original-image level using StratifiedGroupKFold. Each original image and its eight corresponding augmented variants were treated as a single group and assigned to only one fold. Thus, no augmented version of an image assigned to a validation or test partition could appear in the corresponding training partition. This grouping prevents leakage arising from augmentation while preserving the original class distribution through stratification.

However, this grouping provides image-level rather than verified subject-level independence. The source dataset does not contain reliable subject identifiers; therefore, distinct photographs of the same child, if present, could potentially occur in different folds. Consequently, the evaluation should be interpreted as measuring generalization to unseen images rather than strictly to unseen children, and is discussed as a limitation in Section 7.

#### 3.4.2. Data Augmentation

To improve data diversity and enhance the robustness and generalization capability of the FER models, we employed a comprehensive data augmentation strategy (45). Because the same augmentation strategy was applied uniformly across emotion categories, augmentation increased the effective training-set size without changing the underlying class proportions. The resulting class imbalance was therefore retained and explicitly addressed during evaluation through stratified sampling and the use of balanced accuracy and macro-F1 in addition to overall accuracy. The ASD-FER dataset consists of 2,779 original facial images. Each image was first resized to 224 × 224 pixels and then subjected to eight augmentation techniques while preserving its original emotion label. These transformations generate realistic variations in facial appearance, pose, illumination, and imaging conditions. As a result, 22,232 augmented images (2,779 ×8) were generated, yielding a total of 25,011 images, including the original images, Table 3 shows the emotion class distribution before and after augmentation. The augmentation techniques are summarized as follows:

- **Horizontal Flip:** Mirrors the facial image to increase viewpoint diversity and reduce sensitivity to left–right facial orientation.
- **Random Rotation:** Rotates each image by a random angle within [−20^◦^, 20^◦^] to improve robustness to head pose variations.
- **Color Jitter:** Randomly adjusts image brightness, contrast, saturation, and intensity to simulate variations in illumination and camera settings.
- **Gaussian Noise:** Adds zero-mean Gaussian noise (*σ* = 15) to improve robustness against sensor noise and low-quality image acquisition.
- **Image Blending:** Blends each image with its horizontally flipped version using a random interpolation weight (*α* ∈ [0.1, 0.3]), encouraging the models to learn more robust global facial representations.
- **Gaussian Blur:** Applies Gaussian smoothing with a randomly selected kernel size to simulate slight camera defocus and motion blur.
- **Random Affine Transformation:** Introduces small geometric deformations by perturbing the image corner locations, improving robustness to subtle facial shape variations.
- **Gradient-based Illumination Augmentation:** Applies a smooth horizontal or vertical illumination gradient to emulate non-uniform lighting conditions commonly encountered in real-world environments.

**Table 3.** Class-wise distribution of the ASD-FER dataset before augmentation and the additional images generated after applying eight augmentation techniques.

| Emotion Class | Before Augmentation | Augmented Images Generated (8×) |
| --- | --- | --- |
| Angry | 112 | 896 |
| Fear | 82 | 656 |
| Happy | 1,717 | 13,736 |
| Neutral | 758 | 6,064 |
| Sad | 51 | 408 |
| Surprise | 59 | 472 |
| **Total** | **2,779** | **22,232** |

These augmentation strategies (Fig. 4) preserve the semantic content of facial expressions while substantially increasing dataset diversity. This enables both the transformer-based models (HybridT, Swin-T, ViT) and the six conventional baselines (custom CNN, SVM, ResNet34, ResNet50, MobileNetV2, and AlexNet) to learn more robust and invariant feature representations.

**Figure 4:**
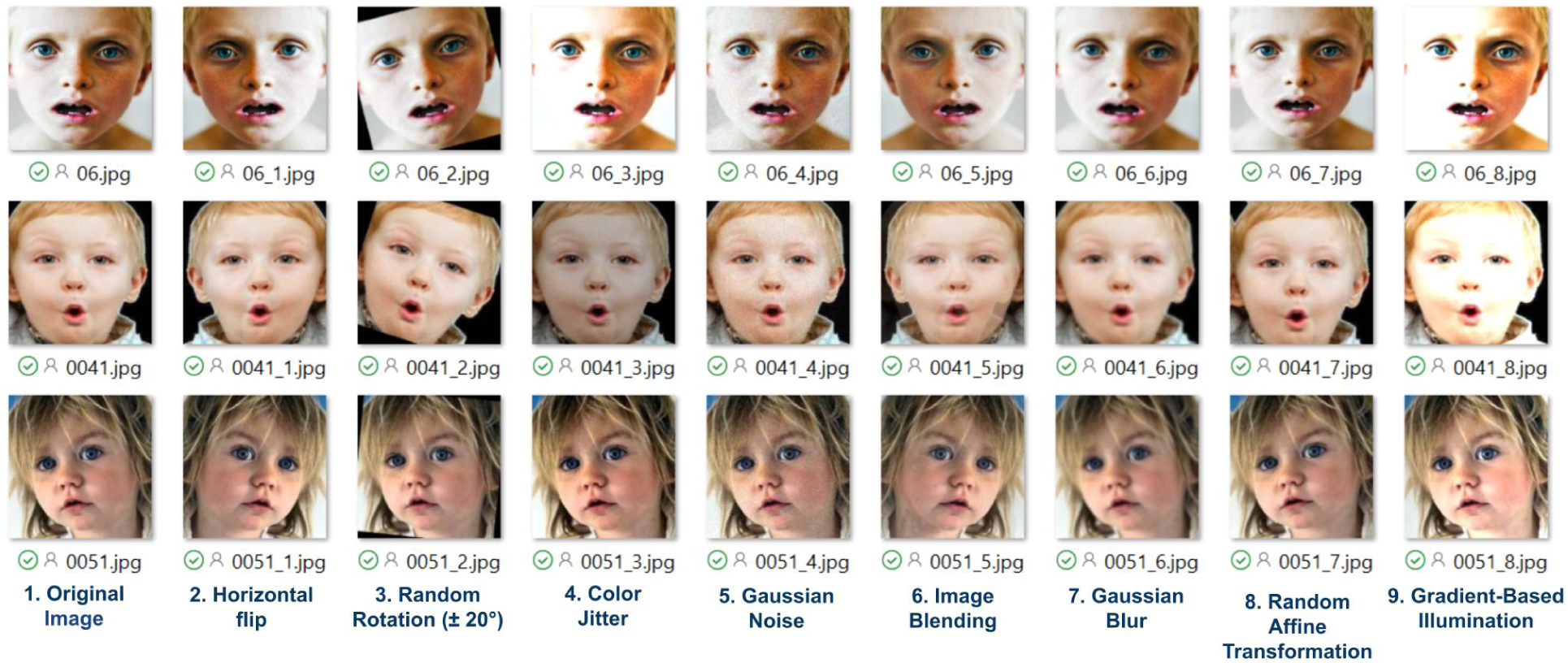
Examples of the eight data augmentation techniques applied to the ASD-FER images. The facial images are from the publicly available Kaggle dataset, Facial_data_ASD by Md. Shafiul Alam (34)

## 4. Experimental Setup

### 4.1. Model Architectures

To establish a comprehensive performance reference, six models spanning classical ML and DL paradigms were benchmarked: a linear-kernel SVM, a custom CNN, and four ImageNet-pretrained architectures (AlexNet, ResNet34, ResNet50, and MobileNetV2). The SVM operates directly on standardized pixel vectors and quantifies the separability of expression classes without any representation learning, while the custom CNN measures what a compact, task-specific network can achieve when trained from scratch. The four pretrained architectures were selected to cover complementary design philosophies: AlexNet as an early deep architecture, ResNet34 and ResNet50 as residual networks of increasing depth, and MobileNetV2 as an efficiency-oriented model representative of resource-constrained deployment. Furthermore, three transformer-based architectures were investigated for ASD facial emotion recognition: a lightweight HybridT, Swin-T, and ViT. These architectures represent three complementary design paradigms: hybrid convolution-transformer feature extraction, hierarchical window-based self-attention, and pure global self-attention.

To enable a controlled comparison among the transformer-based models, all three shared the same downstream architecture, comprising an Emotion Region Enhancement (ERE) module, six learnable Emotion Query Tokens, a two-layer Transformer Decoder, a Contrastive Module, and an Emotion Classifier. These models differ only in their backbone feature extractors, enabling a controlled comparison of the three transformer paradigms. We refer to this shared downstream design – the ERE module, gated Emotion Query Tokens, Transformer Decoder, and dual contrastive/classification heads – as **EQFormer**. Analogous to how DETR (46) pairs a novel query-based decoder with a standard ResNet backbone, the backbone itself is not our contribution; EQFormer is the task-specific architecture built around it, and its ability to generalize is demonstrated here across three different backbones (HybridT, Swin-T, and ViT) evaluated on ASD-FER. Query-token decoders of this kind have recently been explored for unifying multiple facial-analysis tasks — FaceXFormer (18), QFace, and Faceptor each learn one token per task (e.g., parsing, landmarks, age) that cross-attends to a shared backbone representation, unifying up to ten heterogeneous tasks on a single backbone. EQFormer applies the same query-token principle along the orthogonal axis: six gated tokens, one per emotion class, within a single task (FER), while holding the decoder fixed and varying the backbone (HybridT, Swin-T, ViT). This design is what makes EQFormer suited to isolating backbone effects for ASD FER specifically, rather than unifying tasks.

#### 4.1.1. Custom CNN

A custom convolutional neural network (CNN) was implemented as a baseline, consisting of two convolutional layers (16 and 32 filters, 3×3 kernels) with ReLU activations and max pooling, followed by two fully connected layers. CNNs learn hierarchical spatial features directly from pixel data through local receptive fields and weight sharing, making them well suited to FER (47). Its shallow two-block design provides a fast reference point against the deeper pretrained architectures, although the large flattened feature map places most of its parameters in the first fully connected layer.

#### 4.1.2. SVM

The support vector machine (SVM) serves as a classical ML baseline operating on flattened, standardized pixel intensities. A linear-kernel SVM with regularization parameter *C*=1 finds the maximum-margin hyperplane separating classes in the high-dimensional input space (48). Unlike the deep models, it performs no representation learning, so its performance indicates how much of the task is solvable from raw pixel statistics alone.

#### 4.1.3. AlexNet

AlexNet is an eight-layer architecture comprising five convolutional and three fully connected layers, which popularized ReLU activations, dropout regularization, and GPU-based training of deep networks (49). In this study, an ImageNet-pretrained AlexNet was adapted to single-channel input by summing its first-layer kernels across the color dimension, its classification layer was replaced, and the network was fine-tuned using the two-stage regime described below. Under this protocol it collapsed to a degenerate majority-class solution rather than learning discriminative features.

#### 4.1.4. ResNet34

ResNet34 is a 34-layer residual network in which identity shortcut connections allow gradients to propagate directly through the network, mitigating the vanishing-gradient and degradation problems of deep architectures (50). Its basic residual blocks with 3×3 convolutions offer a favorable balance between depth and computational cost. The ImageNet-pretrained model was adapted to grayscale input and fine-tuned in a two-stage regime (five-epoch head warm-up at lr=10^−2^, followed by full fine-tuning at lr=10^−4^, early stopping on validation loss with patience 50).

#### 4.1.5. ResNet50

ResNet50 extends the residual learning framework to 50 layers using bottleneck blocks (1×1, 3×3, 1×1 convolutions) that reduce computation while increasing representational capacity (50). Its greater depth enables richer feature hierarchies than ResNet34 at a modest parameter cost, but it is the most expensive model to train (8,501 s across ten folds against 5,392 s for ResNet34) and has the lowest inference throughput of the deep models. The same transfer learning and two-stage fine-tuning protocol was applied.

#### 4.1.6. MobileNetV2

MobileNetV2 is an efficiency-oriented architecture built on depthwise separable convolutions, inverted residual blocks, and linear bottlenecks, which drastically reduce parameter count and multiply–accumulate operations relative to standard CNNs (51). It represents the resource-constrained deployment scenario relevant to edge and mobile inference, and was fine-tuned from ImageNet weights using the same adaptation procedure as the other pretrained models.

The architectures of the six conventional baselines are summarized in Table 4. All operate on single-channel 224×224 images at the native resolution of the ASD corpus. The linear SVM operates directly on 50,176-dimensional raw pixel features and serves as a non-deep-learning reference, while the custom CNN (12.85M parameters, most of them in its first fully connected layer) is trained using stacked convolutional blocks. The remaining four models–AlexNet (57.01M), ResNet34 (21.28M), ResNet50 (23.51M), and MobileNetV2 (2.23M)–are initialized with ImageNet-1K weights and further pretrained for 30 epochs on the combined FERAC (17) and FER-Autism (33) corpora (1,814 images) before fine-tuning on the ASD dataset. None employs any attention mechanism, providing a clear architectural contrast to the transformer-based models introduced next.

**Table 4.** Comparison of the model architectures evaluated in this study.

| Model | Pretraining | Parameters | Feature Extraction | Attention Type |
| --- | --- | --- | --- | --- |
| Custom CNN | None (trained from scratch) | 12.85M | Stacked Convolutional Blocks | None |
| SVM (linear, $C=1$) | None | N/A<sup>a</sup> | Raw Pixel Features (50,176-d) | None |
| AlexNet | ImageNet-1K → FERAC + FER-Autism | 57.01M | Sequential Convolutions | None |
| ResNet34 | ImageNet-1K → FERAC + FER-Autism | 21.28M | Residual Convolutional Blocks | None |
| ResNet50 | ImageNet-1K → FERAC + FER-Autism | 23.51M | Bottleneck Residual Blocks | None |
| MobileNetV2 | ImageNet-1K → FERAC + FER-Autism | 2.23M | Depthwise Separable Convolutions | None |
| HybridT | FERAC + FER-Autism | 2.66M | CNN + Transformer Encoder | Local + Global |
| Swin-T | ImageNet-1K | 44.7M | Hierarchical Transformer | Shifted-window |
| ViT | ImageNet-21K | 102.95M | Patch Embedding | Global Self-Attention |

<sup>a</sup>The SVM has no learned weight tensor comparable to a network's parameter count; the fitted models retain 1,575–2,115 support vectors.

#### 4.1.7. Hybrid CNN-Transformer

The proposed **HybridT** is a lightweight architecture designed to efficiently capture both local facial characteristics and global contextual relationships for facial emotion recognition in children with ASD. The HybridT model is a compact, base architecture that processes each face image through six sequential stages before splitting into two task-specific output heads. The complete pipeline is illustrated in Fig. 5. The proposed model first extracts low-level visual representations using convolution layers before modeling long-range dependencies through a transformer encoder. This design significantly reduces computational complexity while preserving discriminative facial features. The implementation contains approximately 2.66 million trainable parameters, making it substantially more compact than the transformer backbones evaluated in this study.

a. **Input**: A 224×224×3 aligned face crop enters the pipeline.
b. **Emotion Region Enhancement (ERE)**: A module is first applied to the input image to emphasize facial regions that are most informative for emotion recognition. The module consists of a lightweight convolutional attention block with a 3→16→3 channel structure followed by a sigmoid activation to generate a spatial attention map. The input image is multiplied by this attention map, enabling the network to focus on emotion-relevant facial regions before feature extraction.
c. **CNNStem**: The enhanced image is then processed by a CNNStem, composed of two convolutional layers with Batch Normalization and ReLU activation. The first convolution expands the input from 3 to 64 channels using stride-2 downsampling, while the second increases the feature dimension to 128 channels. These convolutional layers efficiently capture local facial textures and edge information while progressively reducing the spatial resolution.
d. **Transformer Encoder**: The 128-channel feature map is adaptively pooled to a 9×9 grid (81 tokens) and passed through a 2-layer transformer encoder (2 attention heads) that lets every spatial location attend to every other location, building global context on top of the CNN’s local features.
e. **Emotion Query Tokens**: Six learnable query vectors — one per emotion class — are conditioned on the image via a gated projection (Linear → Sigmoid gate), so each query is adapted to the specific face rather than being a fixed, generic template.
f. **Transformer decoder**: A 2-layer decoder lets the six emotion queries cross-attend to the encoder’s 81 spatial tokens, allows each query token to attend selectively to emotion-relevant facial features, producing discriminative representations for the six emotion categories.
g. **Two output heads**: The decoded, class-conditioned representations feed two parallel heads:

- **Contrastive module** — normalizes and passes the pooled representation through a small MLP projection head (128→128→64) into an embedding space used only during training, where a supervised contrastive loss pulls same-emotion samples together and pushes different emotions apart.
- **Emotion classifier** — composed of Layer Normalization, a fully connected layer with Gaussian Error Linear Unit (GELU) activation, dropout regularization, and a final linear classification layer predicts the six facial emotion classes.

**Figure 5:**
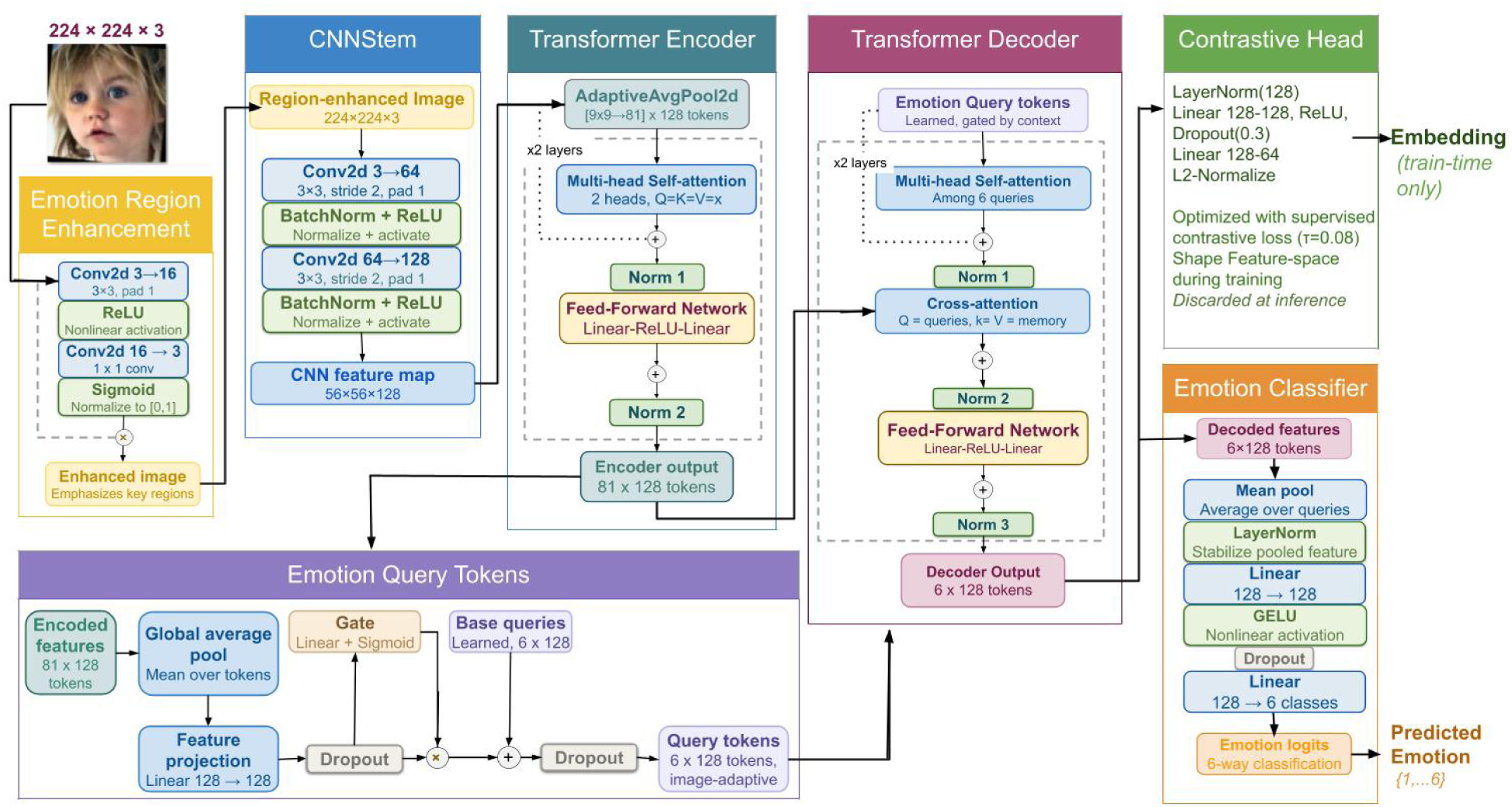
End-to-end architecture of **EQFormer** with the HybridT backbone configuration, from input face image through Emotion Region Enhancement, CNN stem, Transformer Encoder, gated Emotion Query Tokens, and Transformer Decoder, to the training-only Contrastive Head and the inference-time Emotion Classifier. The input facial image is derived from the publicly available Kaggle dataset, Facial_data_ASD, by Md. Shafiul Alam (34).

At inference time, only the classifier branch is used to produce the predicted emotion; the contrastive embedding exists purely to shape a better-separated feature space during training. This whole pipeline totals 2.66M parameters — roughly 17× smaller than Swin-T and 39× smaller than ViT — while still reaching 89.57% held-out test accuracy.

The complete layer-wise architecture summary, including per-layer output shapes and parameter counts, is provided in Appendix B (Fig. 15).

#### 4.1.8. Swin Transformer

The proposed Swin Transformer model builds upon the official Swin-Tiny architecture pretrained on ImageNet-1K, extended with task-specific modules for facial emotion recognition. As illustrated in Fig. 6 — the shared pipeline for both the Swin-T and ViT variants — the model consists of an Emotion Region Enhancement module, the pretrained Swin-T backbone, a query-conditioned decoding stage, and dual classification/contrastive heads. The full layer-wise summary is provided in Appendix B (Fig. 16).

a. **Emotion Region Enhancement**. Same as HybridT — the 3→16→3 convolutional spatial-attention gate applied to the input image before the backbone.
b. **Swin-T backbone (pretrained, new)**. Replaces the HybridT’s CNNStem + custom TransformerEncoder. The enhanced facial image is processed by a pretrained Swin-Tiny backbone (timm library from Hugging Face) initialized with ImageNet-1K weights and fine-tuned end-to-end at a substantially lower learning rate (1e-6) than the downstream modules (5e-5). The backbone first partitions the input into non-overlapping 4×4 patches and projects them into a 96-dimensional embedding space. It then extracts hierarchical features through four stages with embedding dimensions increasing from 96 to 768 and attention heads of [3,6,12,24]. Alternating window-based and shifted-window self-attention effectively capture both local facial details and global contextual information, while patch merging generates multi-scale feature representations. The final backbone features are globally averaged to obtain a compact 768-dimensional feature vector for the subsequent emotion decoding and classification modules.
c. **Token reshaping**. The final 768-dimensional feature vector extracted by the Swin-T backbone is reshaped into a sequence of length one before being forwarded to the shared emotion decoding stage. Unlike the HybridT and ViT, which preserve multiple spatial or patch-level tokens for decoding, the Swin-T backbone provides a single global feature representation. This design allows the subsequent emotion-aware decoder to transform the global embedding into discriminative emotion-specific features while maintaining a unified decoding architecture across all transformer backbones.
d. **Emotion Query Tokens + Decoder**. The decoder architecture is identical to that of the HybridT model, employing six gated learnable Emotion Query Tokens and a two-layer Transformer Decoder, with the embedding dimension increased from 128 to 768. Since the encoder memory consists of a single global feature representation, the decoder does not perform spatial reasoning over multiple tokens. Instead, each learnable emotion query interacts with the shared global representation to generate an emotion-specific feature embedding. Maintaining the same decoding architecture across the HybridT, Swin-T, and ViT models ensures that the backbone feature extractor remains the only architectural difference, enabling a controlled assessment of the different backbone paradigms within the proposed framework.
e. Contrastive module + classifier. Identical design to HybridT, dimensions scaled to 768→128→64 (contrastive) and 768→128→6 (classifier).

**Figure 6:**
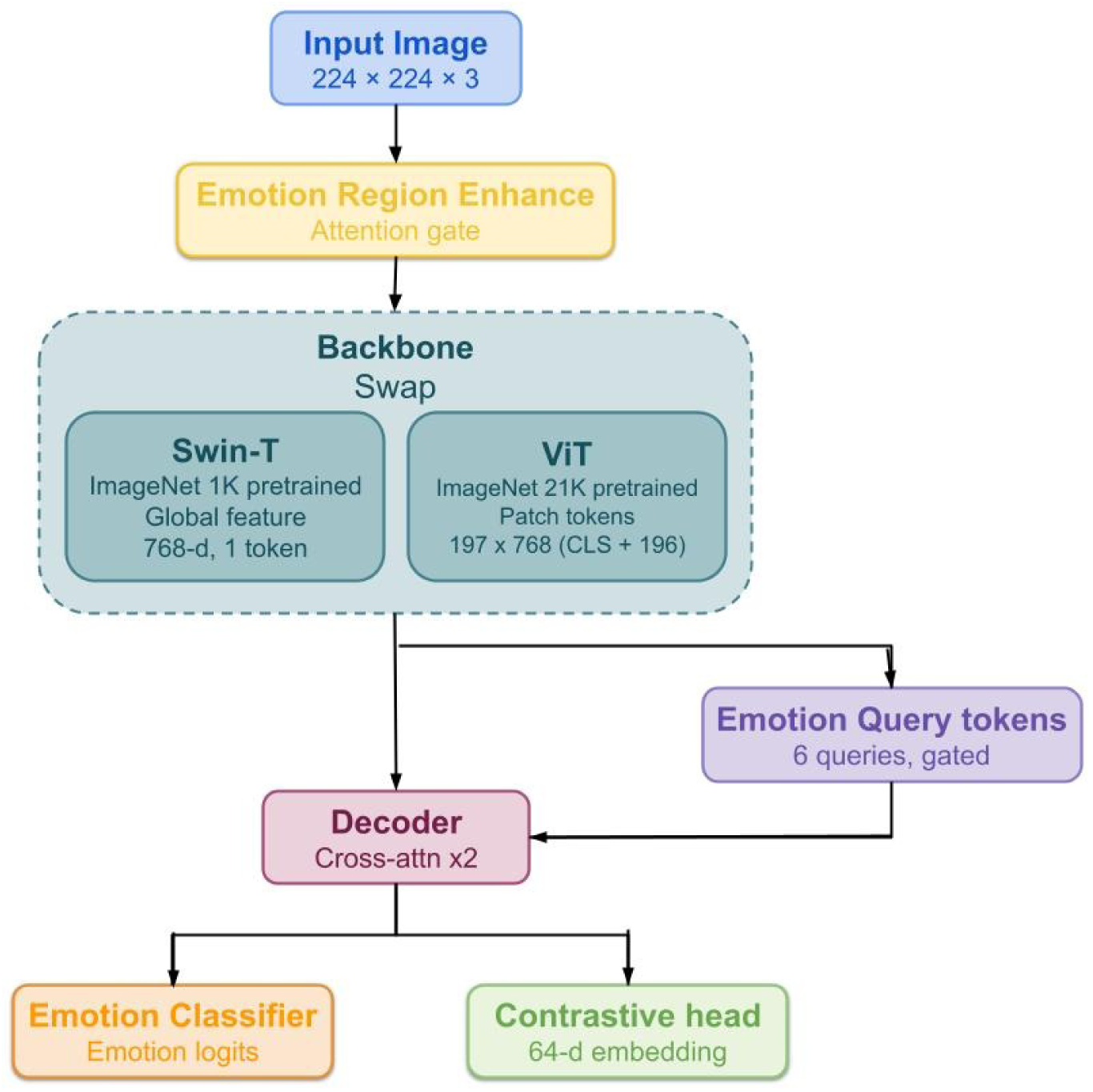
EQFormer with Swin-T and ViT backbone configurations, sharing one downstream pipeline, differing only in the backbone: Swin-T (ImageNet-1K) pools to a single 768-d token, while ViT (ImageNet-21K) preserves all 197 patch tokens at 768-d, giving the decoder richer spatial context to cross-attend over.

Overall, the proposed Swin-T model combines the hierarchical representation capability of shifted-window self-attention with task-specific emotion-aware decoding. By integrating Emotion Region Enhancement, learnable Emotion Query Tokens, supervised contrastive learning, and ImageNet-1K pretraining within a unified architecture, the model effectively captures both local facial characteristics and high-level semantic representations while preserving a controlled architectural comparison with the HybridT and ViT.

#### 4.1.9. Vision Transformer

The ViT was implemented using the official Hugging Face google/vit-base-patch16-224-in21k model pretrained on ImageNet-21K. The input image is partitioned into fixed-size 16 × 16 patches, which are linearly embedded into tokens. A learnable class token and positional embeddings are added before the sequence is processed through twelve Transformer encoder layers using global multi-head self-attention. Similar to Swin-T, an Emotion Region Enhancement module is incorporated before the backbone to emphasize discriminative facial regions. The backbone features are then processed using the same Emotion Query Tokens, Transformer Decoder, contrastive embedding module, and emotion classifier shown in the combined architecture diagram (Fig. 6). Because the ViT backbone retains its full output sequence rather than a single pooled vector, the decoder’s cross-attention operates over 197 spatial/semantic tokens - compared to the single globally-pooled token available in the Swin-T variant - giving the decoder stage substantially richer context to draw from despite reusing an architecturally identical module. Unlike Swin-T, ViT performs global attention over all image patches without a hierarchical feature pyramid, enabling richer global context while sacrificing explicit multi-scale representation. The full layer-wise summary is provided in Appendix B (Fig. 17).

#### 4.1.10. Comparison of Transformer Architectures

The three transformer-based models represent different trade-offs between model complexity and representational capability. The HybridT emphasizes computational efficiency with only 2.66M parameters, whereas Swin-T employs hierarchical shifted-window attention with approximately 44.7M parameters to better capture local and global facial characteristics. ViT represents the largest model (102.95M parameters) and relies entirely on global self-attention with ImageNet-21K pretraining. Evaluating these architectures under an identical experimental protocol enables a controlled comparison of parameter efficiency, transfer learning effectiveness, and recognition performance for ASD facial emotion recognition. The comparison is shown in Table 4.

### 4.2. Training Configuration

All six models (custom CNN, SVM, AlexNet, ResNet34, ResNet50, and MobileNetV2) were evaluated on grayscale images at their native 224×224 resolution with a mini-batch size of 32. The deep learning models were trained using the Adam optimiser with cross-entropy loss on GPU hardware. Pretraining on the combined FER-Autism (33) and FERAC (17) corpus (1,814 images: 1,199 from FER-Autism and 615 from FERAC) was performed for 30 epochs with a learning rate of 10^−3^ and no augmentation. Fine-tuning on the ASD dataset followed a two-stage regime for the pretrained architectures: first, a five-epoch head warm-up in which the backbone was frozen and only the newly initialised classification layer was trained with a learning rate of 10^−2^; second, full end-to-end fine-tuning of all layers at a reduced learning rate of 10^−4^. The custom CNN was trained at a learning rate of 10^−3^. All deep models were fine-tuned for a maximum of 200 epochs with early stopping monitored on the validation loss with a patience of 50 epochs, after which the weights corresponding to the lowest validation loss were restored. The SVM baseline used a linear kernel with regularisation parameter *C*=1, operating on flattened 50,176-dimensional raw pixel features standardised to zero mean and unit variance; it was trained on augmented images during cross-validation and on unaugmented images in the final held-out evaluation. Training hyperparameters are summarised in Table 5.

**Table 5.**
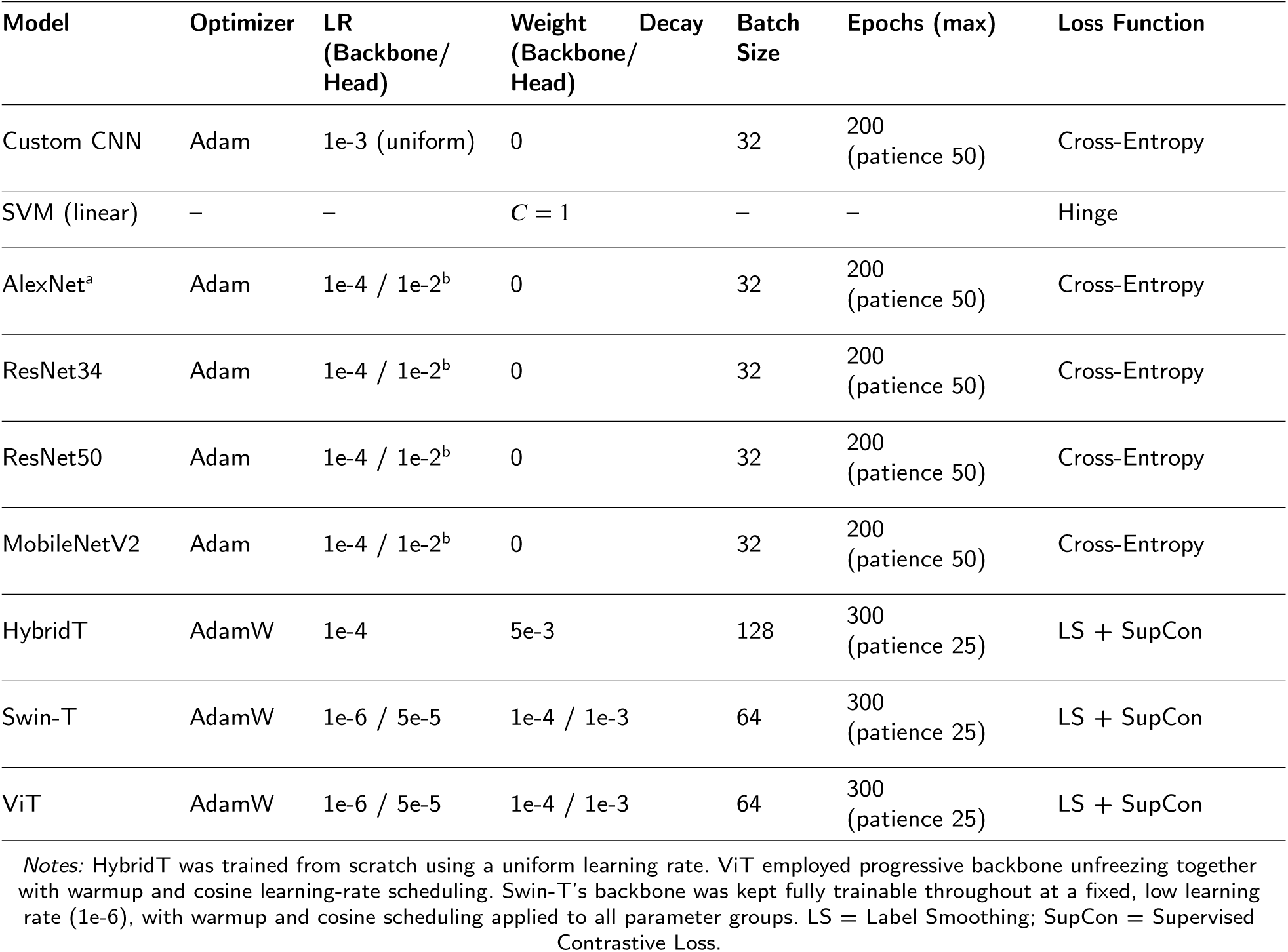
Training hyperparameters used for each model.

All transformer-based models were optimized using the AdamW optimizer with cosine annealing learning-rate scheduling. For the pretrained Swin-T and ViT models, differential learning rates were employed by assigning a smaller learning rate to the backbone and a larger learning rate to the task-specific modules, allowing effective adaptation while preserving pretrained representations. In contrast, the HybridT — which does not use ImageNet-pretrained weights — used a single learning rate for all network parameters. The HybridT model instead followed a two-stage training strategy of its own: domain-specific pretraining on the FERAC (17) and FER-Autism (33) datasets, followed by fine-tuning on the ASD-FER dataset, whereas Swin-T and ViT were directly fine-tuned from their ImageNet-pretrained weights.

To improve both classification performance and feature discrimination, all models were trained using a joint objective combining label-smoothed cross-entropy loss with supervised contrastive loss. The label smoothing factor was set to 0.03 for the HybridT model and 0.02 for Swin-T and ViT. Batch size, training epochs, learning rates, weight decay, and other hyperparameters (52) are summarized in Table 5. All experiments were conducted using the PyTorch deep learning framework on a workstation equipped with an NVIDIA GeForce RTX 5070 Ti GPU (16 GB VRAM).

A progressive backbone unfreezing strategy was applied to the pretrained ViT model, where the encoder layers were gradually unfrozen during training to facilitate stable adaptation while retaining pretrained knowledge. For Swin-T, the pretrained backbone was instead kept fully trainable throughout training at a fixed, very small learning rate (1e-6), relying on the differential learning-rate scheme in Table 5 to balance backbone stability against task-specific adaptation. Since the HybridT does not rely on ImageNet-pretrained weights, all network parameters remained trainable throughout both the pretraining and fine-tuning stages.

### 4.3. Evaluation Protocols

All models were evaluated under two protocols. First, stratified 10-fold cross-validation with an 80/10/10 train-validation-test split was used to fine-tune each model on ASD-FER via transfer learning, ensuring robust and reproducible performance estimates across all folds. Second, a final held-out evaluation on a 90/10 split provided an unbiased assessment of generalization to unseen data.

#### 4.3.1. Stratified 10-Fold Cross-Validation

Model performance was assessed using stratified 10-fold cross-validation applied to the pool of 2,779 original images. StratifiedGroupKFold (random_state=42, see Fig. 2) was applied to the original images only, preserving the class distribution of the six emotion (*Angry, Fear, Happy, Neutral, Sad, Surprise*) in every fold. For each fold, the data were partitioned as follows (see Fig. 7):

- **Training set (80%, 8 folds):** original images plus their corresponding augmented variants (from a pool of 22,232 augmented images), trained with a full random augmentation pipeline.
- **Validation set (10%, 1 fold):** original images plus corresponding augmented variants.
- **Test set (10%, 1 fold):** original images only, clean and untouched (no augmentation applied).

**Figure 7:**
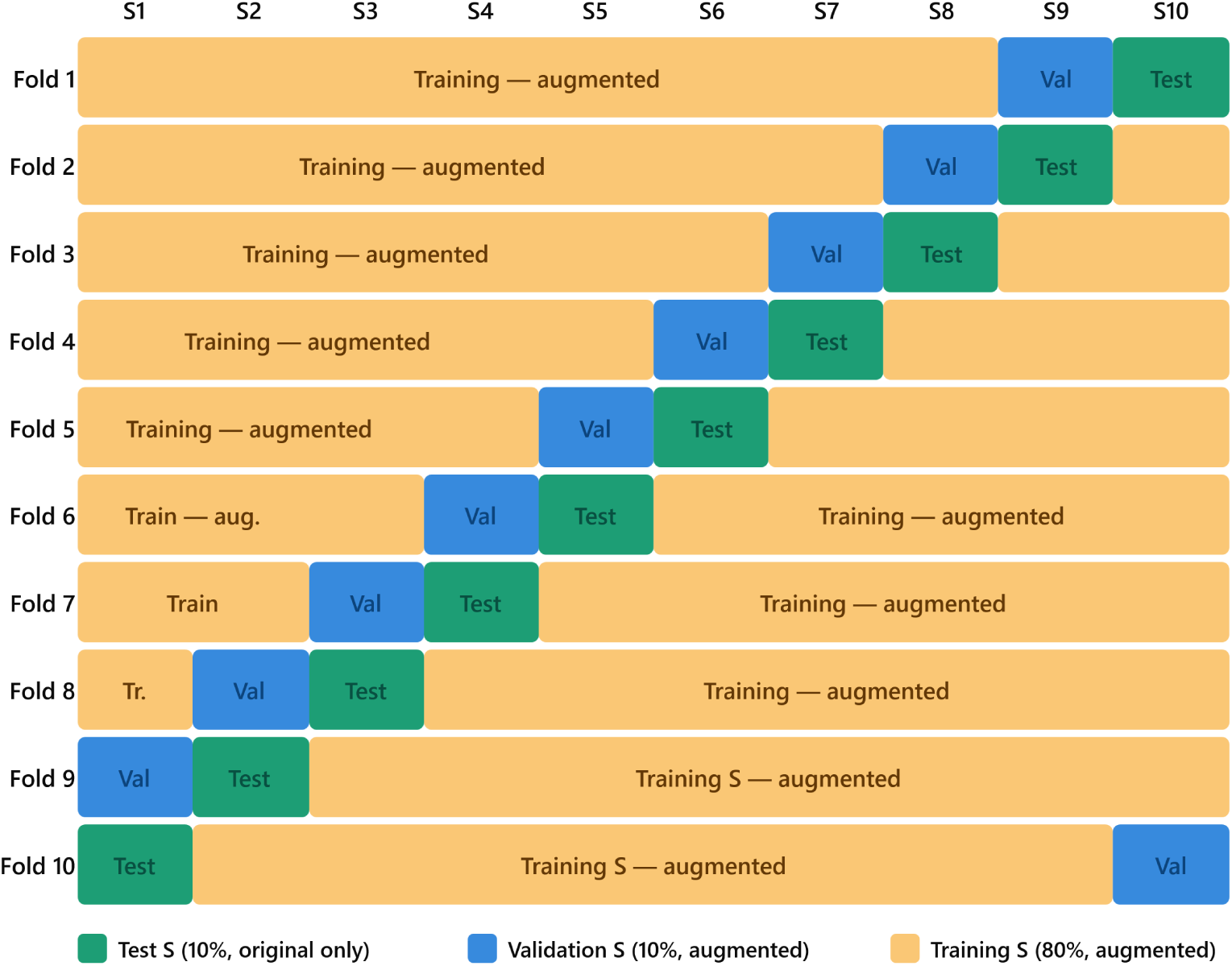
Stratified 10-fold cross-validation scheme. Each fold assigns one subset as the test set (10%), one as the validation set (10%), and the remaining eight as the training set (80%). All non-test splits are augmented. S = subset; folds rotate systematically so every subset serves as test exactly once.

This design ensures that augmentation-derived variants of a given source image never leak across the train/validation/test boundary within a fold, while still providing an augmented training signal. Performance was measured using six metrics: accuracy, balanced accuracy, precision, recall, weighted F1, and macro-F1, each computed per fold and averaged across all 10 folds for both the validation and test partitions. Balanced accuracy and macro-F1 were emphasized as the primary metrics for comparing model robustness on minority emotion classes (*Fear, Sad, Surprise*), given the pronounced class imbalance in the dataset.

#### 4.3.2. Held-Out Final Evaluation

Following cross-validation, a final model was trained on the full available dataset (original training images merged with the complete augmented image pool), using a 90/10 train/test split (train_test_split, stratified by class, random_state=42) rather than the fold-based partitioning used during cross-validation. As with the cross-validation protocol, the split was performed prior to augmentation: only the 90% training partition was expanded with its augmented variants, while the 10% test partition retained only original, unaugmented images, so that final performance is measured exclusively on clean held-out images. This final model was initialized from the same pretrained checkpoint and trained under the same optimizer/scheduler/loss configuration as the per-fold models (Table 5), tracked with early stopping on validation loss.

The purpose of this held-out evaluation is to obtain a single, unbiased estimate of the model’s generalization performance using its maximum available training data — beyond what any single cross-validation fold can offer — and to produce the final deployable model checkpoint. The same six metrics (accuracy, balanced accuracy, precision, recall, weighted F1, macro-F1) were computed on the held-out test partition, along with a full per-class classification report (precision/recall/F1 per emotion) to characterize class-wise generalization.

### 4.4. Evaluation Metrics

Model performance was evaluated using overall accuracy, balanced accuracy, precision, recall, weighted F1-score, and macro F1-score. Overall accuracy measures the proportion of correctly classified samples, whereas balanced accuracy accounts for class imbalance by averaging recall across all classes. Precision and recall evaluate the reliability and completeness of predictions, respectively. Weighted F1-score reflects class performance while considering class frequencies, whereas macro F1-score assigns equal importance to each class regardless of its size. Table 6 presents the performance measurement metrics (53) used to evaluate the multi-class FER task.

**Table 6.** Evaluation metrics used for facial expression recognition (FER) performance assessment.

| Metric | Function | Formula | Eq. No. |
| --- | --- | --- | --- |
| Accuracy | Presents the overall proportion of correctly classified facial expressions across all six emotion classes. | $\frac{\sum_{i=1}^K TP_i}{N}$ | (4.1) |
| Balanced Accuracy | Computes the average per-class recall across all $K$ emotion classes, giving equal weight to majority and minority classes regardless of their sample size. | $\frac{1}{K} \sum_{i=1}^K \frac{TP_i}{TP_i + FN_i}$ | (4.2) |
| Precision | Calculates the proportion of correctly recognized samples among all samples predicted as a given emotion class, averaged across classes and weighted by class support. | $\sum_{i=1}^K \frac{n_i}{N} \cdot \frac{TP_i}{TP_i + FP_i}$ | (4.3) |
| Recall | Proportion of correctly recognized samples among all true samples of a given emotion class, averaged across classes and weighted by class support. | $\sum_{i=1}^K \frac{n_i}{N} \cdot \frac{TP_i}{TP_i + FN_i}$ | (4.4) |
| Weighted F1 | Combines precision and recall into a single per-class metric, then averages across classes weighted by the number of true samples in each class, reflecting performance at the dataset's natural class distribution. | $\sum_{i=1}^K \frac{n_i}{N} \cdot \frac{2 \cdot Precision_i \cdot Recall_i}{Precision_i + Recall_i}$ | (4.5) |
| Macro F1 | Calculates the F1-score for each emotion class independently and averages them without weighting by class size, treating all six emotion classes equally regardless of sample frequency. | $\frac{1}{K} \sum_{i=1}^K \frac{2 \cdot Precision_i \cdot Recall_i}{Precision_i + Recall_i}$ | (4.6) |

Because the ASD-FER dataset is highly imbalanced, balanced accuracy and macro F1-score were considered the primary evaluation metrics, as they provide a more reliable assessment of model performance on minority emotion classes than overall accuracy alone.

## 5. Results and Discussion

### 5.1. Cross-Validation Results

#### 5.1.1. Conventional and CNN-Based Models

Tables 7 and 8 report fold-wise accuracy and macro F1 for the six conventional and CNN-based baselines under stratified 10-fold cross-validation. The three transfer-learned backbones are closely matched and separate differently under the two metrics: ResNet50 attains the highest mean test accuracy (84.89%) but ResNet34 the highest mean macro F1 (0.5972 against 0.5739), with MobileNetV2 third on both (84.13%, 0.5362). This reversal reflects macro F1’s equal weighting of the six classes, under which ResNet34’s minority-class handling outweighs ResNet50’s majority-class accuracy advantage. All three exceed the from-scratch CNN (78.88%, 0.5066) consistently across folds, although fold-level variability is substantial: ResNet50’s test macro F1 ranges from 0.3620 (F5) to 0.7186 (F0), reflecting the sensitivity of minority-class scores to which handful of rare-class samples land in each 278-image test partition.

**Table 7.** Fold-wise accuracy and macro-F1 on the validation split, stratified 10-fold cross-validation.

| Model | Metric | F0 | F1 | F2 | F3 | F4 | F5 | F6 | F7 | F8 | F9 | Mean |
| --- | --- | --- | --- | --- | --- | --- | --- | --- | --- | --- | --- | --- |
| Custom CNN | Acc | 0.7374 | 0.7626 | 0.7662 | 0.7698 | 0.7158 | 0.7590 | 0.7194 | 0.7590 | 0.7437 | 0.7230 | **0.7456** |
|  | Macro F1 | 0.3601 | 0.4802 | 0.4437 | 0.5081 | 0.2340 | 0.2857 | 0.4142 | 0.2978 | 0.4300 | 0.2949 | **0.3749** |
| SVM | Acc | 0.4892 | 0.5144 | 0.5252 | 0.5036 | 0.5576 | 0.5108 | 0.5324 | 0.5180 | 0.4368 | 0.4532 | **0.5041** |
|  | Macro F1 | 0.1575 | 0.1660 | 0.1764 | 0.1740 | 0.1818 | 0.1851 | 0.1547 | 0.1603 | 0.1554 | 0.1420 | **0.1653** |
| ResNet34 | Acc | 0.8381 | 0.8381 | 0.8741 | 0.8129 | 0.8129 | 0.8345 | 0.8094 | 0.8201 | 0.7906 | 0.8633 | **0.8294** |
|  | Macro F1 | 0.5479 | 0.5391 | 0.6551 | 0.4015 | 0.4757 | 0.5172 | 0.5104 | 0.5693 | 0.4277 | 0.6239 | **0.5268** |
| ResNet50 | Acc | 0.8777 | 0.8345 | 0.8201 | 0.8525 | 0.8058 | 0.7914 | 0.8381 | 0.8058 | 0.8231 | 0.8094 | **0.8258** |
|  | Macro F1 | 0.5978 | 0.5555 | 0.5144 | 0.5577 | 0.3731 | 0.3002 | 0.6066 | 0.4240 | 0.3882 | 0.4957 | **0.4813** |
| MobileNetV2 | Acc | 0.8525 | 0.8201 | 0.8561 | 0.8813 | 0.7770 | 0.8165 | 0.8237 | 0.7842 | 0.8087 | 0.8381 | **0.8258** |
|  | Macro F1 | 0.5713 | 0.5317 | 0.6868 | 0.6409 | 0.3665 | 0.3714 | 0.5008 | 0.3113 | 0.3727 | 0.5546 | **0.4908** |
| AlexNet | Acc | 0.6151 | 0.6151 | 0.6151 | 0.6187 | 0.6187 | 0.6187 | 0.6187 | 0.6187 | 0.6209 | 0.6187 | **0.6178** |
|  | Macro F1 | 0.1269 | 0.1269 | 0.1269 | 0.1274 | 0.1274 | 0.1274 | 0.1274 | 0.1274 | 0.1277 | 0.1274 | **0.1273** |
| HybridT | Acc | 0.8553 | 0.8525 | 0.8557 | 0.8553 | 0.8665 | 0.8553 | 0.8169 | 0.8412 | 0.8708 | 0.8375 | **0.8507** |
|  | Macro F1 | 0.6611 | 0.6453 | 0.5550 | 0.6575 | 0.6323 | 0.7611 | 0.5312 | 0.6314 | 0.6761 | 0.6472 | **0.6398** |
| Swin-T | Acc | 0.8633 | 0.8749 | 0.8829 | 0.8877 | 0.8821 | 0.8841 | 0.8397 | 0.8604 | 0.8925 | 0.8690 | **0.8737** |
|  | Macro F1 | 0.6702 | 0.6906 | 0.5787 | 0.7026 | 0.6656 | 0.7734 | 0.5542 | 0.6448 | 0.7058 | 0.6788 | **0.6665** |
| ViT | Acc | 0.8677 | 0.8881 | 0.8801 | 0.8877 | 0.8801 | 0.8821 | 0.8401 | 0.8660 | 0.8941 | 0.8662 | **0.8752** |
|  | Macro F1 | 0.6823 | 0.6936 | 0.5607 | 0.7123 | 0.6509 | 0.7729 | 0.5505 | 0.6469 | 0.6837 | 0.7070 | **0.6661** |

**Table 8.** Fold-wise accuracy and macro-F1 on the test split, stratified 10-fold cross-validation.

| Model | Metric | F0 | F1 | F2 | F3 | F4 | F5 | F6 | F7 | F8 | F9 | Mean |
| --- | --- | --- | --- | --- | --- | --- | --- | --- | --- | --- | --- | --- |
| Custom CNN | Acc | 0.8201 | 0.7842 | 0.8237 | 0.7842 | 0.7482 | 0.7518 | 0.8058 | 0.7698 | 0.8273 | 0.7726 | **0.7888** |
|  | Macro F1 | 0.5661 | 0.5234 | 0.5979 | 0.4879 | 0.4437 | 0.3579 | 0.5480 | 0.4765 | 0.6052 | 0.4592 | **0.5066** |
| SVM | Acc | 0.4892 | 0.5324 | 0.5324 | 0.3633 | 0.4676 | 0.4065 | 0.4856 | 0.3417 | 0.5036 | 0.3610 | **0.4483** |
|  | Macro F1 | 0.1602 | 0.1606 | 0.1709 | 0.1668 | 0.1377 | 0.1812 | 0.1572 | 0.1477 | 0.1876 | 0.1272 | **0.1597** |
| ResNet34 | Acc | 0.8669 | 0.8777 | 0.8489 | 0.8453 | 0.8633 | 0.8022 | 0.8633 | 0.8129 | 0.8094 | 0.8773 | **0.8467** |
|  | Macro F1 | 0.6492 | 0.6882 | 0.5981 | 0.5673 | 0.6682 | 0.4680 | 0.6508 | 0.5753 | 0.4621 | 0.6444 | **0.5972** |
| ResNet50 | Acc | 0.8921 | 0.8633 | 0.8741 | 0.8489 | 0.8669 | 0.7806 | 0.8561 | 0.8237 | 0.8345 | 0.8484 | **0.8489** |
|  | Macro F1 | 0.7186 | 0.6429 | 0.6035 | 0.5584 | 0.6299 | 0.3620 | 0.5739 | 0.5529 | 0.5437 | 0.5533 | **0.5739** |
| MobileNetV2 | Acc | 0.8705 | 0.8201 | 0.8525 | 0.8993 | 0.8345 | 0.7770 | 0.8453 | 0.8022 | 0.8525 | 0.8592 | **0.8413** |
|  | Macro F1 | 0.6762 | 0.5264 | 0.6270 | 0.7474 | 0.4867 | 0.3682 | 0.6163 | 0.3936 | 0.4416 | 0.4783 | **0.5362** |
| AlexNet | Acc | 0.6187 | 0.6151 | 0.6151 | 0.6151 | 0.6187 | 0.6187 | 0.6187 | 0.6187 | 0.6187 | 0.6209 | **0.6178** |
|  | Macro F1 | 0.1274 | 0.1269 | 0.1269 | 0.1269 | 0.1274 | 0.1274 | 0.1274 | 0.1274 | 0.1274 | 0.1277 | **0.1273** |
| HybridT | Acc | 0.8495 | 0.8705 | 0.8633 | 0.8633 | 0.8777 | 0.8957 | 0.8741 | 0.8094 | 0.8700 | 0.8845 | **0.8658** |
|  | Macro F1 | 0.6419 | 0.6585 | 0.6663 | 0.6059 | 0.6719 | 0.6968 | 0.7591 | 0.5118 | 0.7364 | 0.6789 | **0.6627** |
| Swin-T | Acc | 0.8638 | 0.8777 | 0.8849 | 0.8813 | 0.8921 | 0.8921 | 0.8921 | 0.8345 | 0.8881 | 0.8953 | **0.8802** |
|  | Macro F1 | 0.6735 | 0.6651 | 0.6943 | 0.5950 | 0.6838 | 0.6877 | 0.7913 | 0.5503 | 0.7419 | 0.7003 | **0.6783** |
| ViT | Acc | 0.8781 | 0.8885 | 0.8921 | 0.8741 | 0.8885 | 0.8813 | 0.9065 | 0.8273 | 0.8953 | 0.8953 | **0.8827** |
|  | Macro F1 | 0.6789 | 0.6976 | 0.7004 | 0.6166 | 0.6765 | 0.6740 | 0.8054 | 0.5259 | 0.7411 | 0.6894 | **0.6806** |

The pixel-based linear SVM performs markedly worse than all deep models (44.83% mean test accuracy, 0.1597 macro F1), confirming that raw-pixel representations are insufficient for this task; its predictions are largely confined to the two majority classes, with mean test F1 of 0.59 on *happy* and 0.29 on *neutral* and essentially zero elsewhere. AlexNet fails outright, collapsing to a majority-class predictor in every fold: its accuracy (mean 61.78%) simply mirrors the prevalence of the *happy* class, while its macro F1 of 0.1273 corresponds to non-zero recall on that single class. We attribute this collapse to the combination of AlexNet’s large, weakly regularized fully connected head (57M parameters) and the severe class imbalance, which drives the network into the trivial majority solution early in fine-tuning; the near-zero variance of its metrics across folds (±0.0002 macro F1) indicates a systematic optimization failure rather than split-dependent noise.

The gap between accuracy and macro F1 for all models highlights the impact of class imbalance. Per-class results aggregated over the ten folds show that *happy* (1,717 samples) is by far the easiest class, with mean test F1 between 0.88 (custom CNN) and 0.94 (MobileNetV2), and *neutral* (758 samples) follows at 0.70–0.78. Performance degrades sharply on the minority classes. *Fear* (82 samples) is the most difficult emotion: mean test F1 ranges from only 0.22 (MobileNetV2) to 0.36 (ResNet34), with recall at zero in several individual folds. *Sad* (51 samples) and *surprise* (59 samples) are similarly unstable, with fold-to-fold standard deviations of the same order as the means themselves (e.g., ResNet50 *sad* F1 0.32 ± 0.21, MobileNetV2 *sad* F1 0.30 ± 0.28), since each test fold contains only five to six samples of these classes. *Angry* (112 samples) occupies an intermediate position (F1 0.38–0.55). These patterns indicate that the models learn robust representations for well-supported expressions but struggle to generalize on under-represented ones, motivating the class-balanced evaluation protocol and the comparison with transformer-based architectures in the following section.

#### 5.1.2. Transformer-Based Models

The three transformer-based models— HybridT, Swin-T, and ViT were evaluated using the stratified 10-fold cross-validation protocol. ViT achieved the highest mean test accuracy (88.27%), followed closely by Swin-T (88.02%), while the lightweight HybridT attained 86.58%. The relatively small performance gap (1.69 percentage points between the highest- and lowest-performing models) indicates that all three architectures effectively learn discriminative facial representations despite their different design philosophies and parameter counts (2.66M for HybridT, 44.7M for Swin-T, and 102.95M for ViT).

Table 7 and Table 8 present the fold-wise validation and test results, respectively. Overall, the three models showed stable performance with only modest fold-to-fold variation, demonstrating robustness despite the severe class imbalance of the ASD-FER dataset. Fold 7 consistently produced the lowest test accuracy and macro-F1 for all three models (HybridT: 80.94% / 0.5118, Swin-T: 83.45% / 0.5503, and ViT: 82.73% / 0.5259), suggesting that this split contained a more challenging distribution of minority-class samples. In contrast, Fold 6 yielded the highest macro-F1 for all three models (0.7591, 0.7913, and 0.8054 for HybridT, Swin-T, and ViT, respectively), indicating that differences across folds were influenced more by data composition than by model architecture. The standard deviation of test accuracy was also comparable across models (HybridT: 2.23%, Swin-T: 1.76%, and ViT: 2.05%), with Swin-T exhibiting the most consistent performance.

The observed performance differences can be attributed to the backbone architectures. ViT employs global self-attention to model relationships across all image patches, enabling effective learning of long-range facial dependencies. Swin-T adopts hierarchical shifted-window attention, capturing both local facial details and multi-scale contextual information with greater computational efficiency. In contrast, the HybridT combines convolutional layers for local feature extraction with transformer encoding for global context, achieving competitive performance while requiring substantially fewer parameters. These results demonstrate that lightweight HybridT architectures can provide an effective balance between recognition performance and computational efficiency.

#### 5.1.3. Head-to-Head Comparison

Table 9 summarizes the mean cross-validation performance of all nine evaluated models and supports three observations that the fold-wise tables cannot. On validation, ResNet34 obtained the best score on all six metrics, whereas ResNet50 was the weakest of the three pretrained CNN backbones on the class-balanced measures (0.4662 balanced accuracy, 0.4813 macro-F1) despite matching MobileNetV2 on accuracy (82.58%); the three should therefore be treated as indistinguishable on accuracy, with ResNet34 preferred on class-balanced grounds. The four trained CNN models (custom CNN, ResNet34, ResNet50 and MobileNetV2) all improved from validation to test, but by only 1.6-4.3 accuracy points against 5.8-11.9 points of balanced accuracy, locating the shift in the minority-class scores rather than in the models themselves. Notably, the accuracy-to-balanced-accuracy gap remained large for every baseline, from 0.2594 (ResNet34) to 0.3145 (custom CNN): the strongest conventional model reached a mean per-class recall of only 0.59.

**Table 9.** Stratified 10-fold cross-validation performance across all nine evaluated models (Custom CNN, SVM, ResNet34, ResNet50, MobileNetV2, AlexNet, HybridT, Swin-T, and ViT), reported as the mean validation and test scores across the 10 folds. Metrics include accuracy, balanced accuracy, precision, recall, weighted F1, and macro-F1. Bold values indicate the best result for each evaluation metric.

| Model | Split | Accuracy | Balanced Acc. | Precision | Recall | Weighted F1 | Macro F1 |
| --- | --- | --- | --- | --- | --- | --- | --- |
| Custom CNN | Validation | 0.7456 | 0.3558 | 0.7191 | 0.7456 | 0.7237 | 0.3749 |
|  | Test | 0.7888 | 0.4743 | 0.7826 | 0.7888 | 0.7777 | 0.5066 |
| SVM | Validation | 0.5041 | 0.1686 | 0.4658 | 0.5041 | 0.4828 | 0.1653 |
|  | Test | 0.4483 | 0.1695 | 0.4584 | 0.4483 | 0.4446 | 0.1597 |
| ResNet34 | Validation | 0.8294 | 0.5068 | 0.8164 | 0.8294 | 0.8184 | 0.5268 |
|  | Test | 0.8467 | 0.5873 | 0.8429 | 0.8467 | 0.8392 | 0.5972 |
| ResNet50 | Validation | 0.8258 | 0.4662 | 0.8155 | 0.8258 | 0.8139 | 0.4813 |
|  | Test | 0.8489 | 0.5587 | 0.8374 | 0.8489 | 0.8396 | 0.5739 |
| MobileNetV2 | Validation | 0.8258 | 0.4843 | 0.8082 | 0.8258 | 0.8140 | 0.4908 |
|  | Test | 0.8413 | 0.5422 | 0.8307 | 0.8413 | 0.8338 | 0.5362 |
| AlexNet | Validation | 0.6178 | 0.1667 | 0.3817 | 0.6178 | 0.4719 | 0.1273 |
|  | Test | 0.6178 | 0.1667 | 0.3817 | 0.6178 | 0.4719 | 0.1273 |
| HybridT | Validation | 0.8507 | 0.6462 | 0.8510 | 0.8507 | 0.8474 | 0.6398 |
|  | Test | 0.8658 | 0.6635 | 0.8693 | 0.8658 | 0.8630 | 0.6627 |
| Swin-T | Validation | 0.8737 | 0.6688 | 0.8741 | 0.8737 | 0.8704 | **0.6665** |
|  | Test | 0.8802 | 0.6779 | 0.8810 | 0.8802 | 0.8769 | 0.6783 |
| ViT | Validation | **0.8752** | **0.6694** | **0.8746** | **0.8752** | **0.8710** | 0.6661 |
|  | Test | **0.8827** | **0.6862** | **0.8838** | **0.8827** | **0.8792** | **0.6806** |

Among the transformer-based models, ViT achieved the best overall performance, obtaining the highest mean test accuracy (88.27%), balanced accuracy (0.6862), and macro-F1 (0.6806). Swin-T followed closely with 88.02% accuracy, 0.6779 balanced accuracy, and 0.6783 macro-F1, while the lightweight HybridT achieved 86.58% accuracy, 0.6635 balanced accuracy, and 0.6627 macro-F1. Notably, Swin-T achieved balanced accuracy and macro-F1 within roughly 0.01 of ViT’s while using less than half its parameters (44.7M vs. 102.95M), underscoring the parameter-efficiency trade-off discussed in Section 4.1.

The architectural basis for these differences among the three transformer models is discussed in Section 4.1; compared with the conventional deep learning models, the results here confirm that pretrained transformer backbones provide a consistent advantage for facial emotion recognition in children with ASD, while the HybridT remains an attractive alternative for resource-constrained applications.

### 5.2. Held-Out Final Evaluation Results

Table 10 presents the performance of all nine models on the independent held-out test set. In general, the transformer-based models consistently outperformed the conventional CNN-based and classical ML models across all evaluation metrics. Among the conventional models, ResNet50 was the clear best performer, achieving the highest accuracy (0.8705), balanced accuracy (0.6440), and Macro F1 (0.6440) of the group. The remaining CNN-family models — the custom CNN, ResNet34, and MobileNetV2 — clustered closely together, each reaching an accuracy between 0.8525 and 0.8633 and a Macro F1 between 0.6181 and 0.6363, indicating that no single conventional backbone besides ResNet50 held a decisive advantage on this dataset. The baseline CNN’s competitiveness (0.8525 accuracy, 0.6363 macro F1) is notable given that it lacks ImageNet pretraining, performing comparably to the pretrained ResNet34 and MobileNetV2 architectures. The pixel-based SVM trailed the deep models substantially (0.7806 accuracy, 0.5956 macro F1), showing that learned convolutional features provide a clear benefit over raw-pixel decision boundaries when the full 90% training pool is available. AlexNet performed markedly worse (balanced accuracy 0.1667, macro F1 0.1274), collapsing to majority-class predictions and failing to generalize to minority classes, consistent with its behavior under cross-validation. Fig. 8 illustrates the loss and accuracy graph for five traditional models; the convergence curve of SVM is not shown because it does not produce per-epoch training curves as it optimises a convex quadratic programme in a single pass.

**Figure 8:**
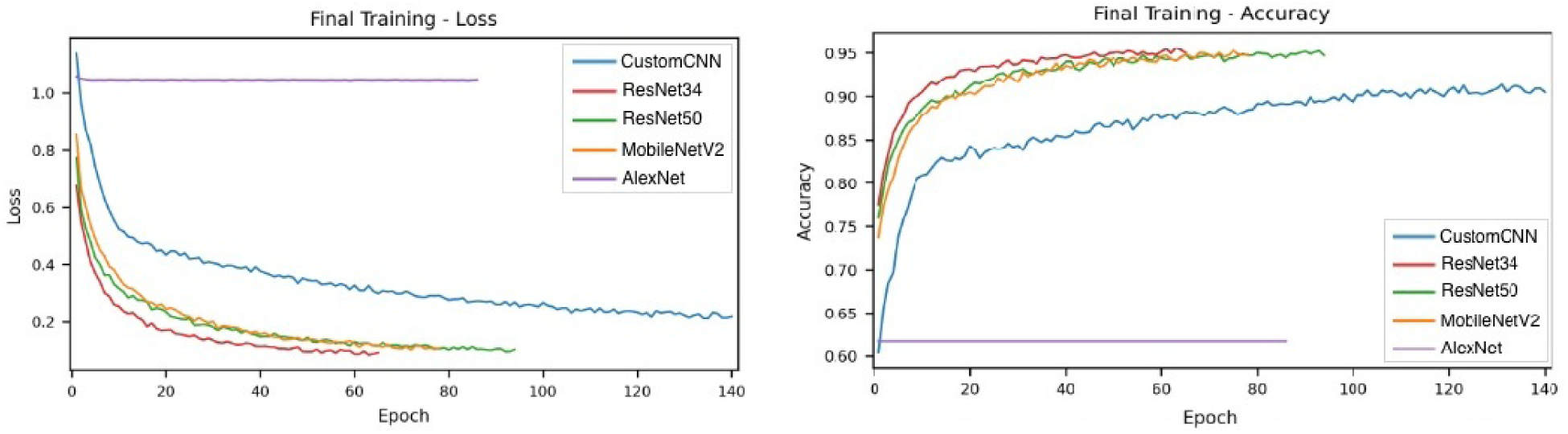
Training loss and accuracy curves for the five traditional models on the 90/10 split

**Figure 9:**
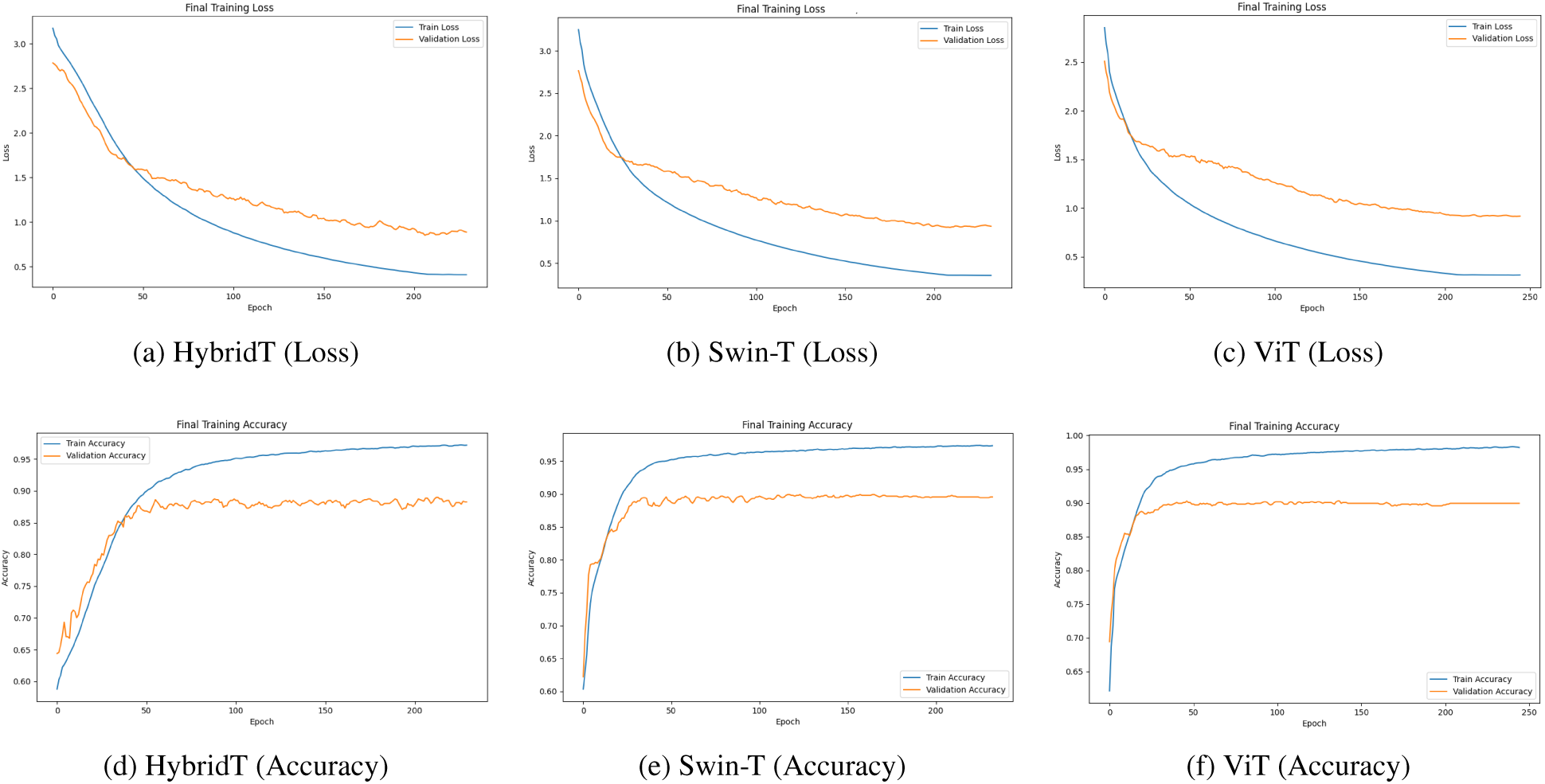
Training loss (top row) and accuracy (bottom row) curves for the HybridT, Swin-T, and ViT models on the 90/10 split. All three models exhibit stable convergence with no evidence of severe overfitting.

**Table 10.** Comparison of held-out (90/10 split) final evaluation performance across all evaluated models, including conventional CNN-based and transformer-based architectures. Bold values indicate the best result for each metric.

| Model | Accuracy | Balanced Accuracy | Macro F1 |
| --- | --- | --- | --- |
| Custom CNN | 0.8525 | 0.6107 | 0.6363 |
| SVM | 0.7806 | 0.5791 | 0.5956 |
| ResNet34 | 0.8633 | 0.6408 | 0.6360 |
| ResNet50 | **0.8705** | **0.6440** | **0.6440** |
| MobileNetV2 | 0.8633 | 0.6383 | 0.6181 |
| AlexNet | 0.6187 | 0.1667 | 0.1274 |
| HybridT | 0.8957 | 0.7301 | 0.6924 |
| Swin-T | 0.9029 | **0.7447** | **0.7078** |
| ViT | **0.9101** | 0.7352 | 0.7024 |

The three transformer-based models achieved the strongest overall results, each exceeding the best conventional models (ResNet50, 87.05% accuracy, 0.6440 macro-F1) by a clear margin. ViT obtained the highest accuracy of 91.01%, while Swin-T achieved the best balanced accuracy (0.7447) and macro-F1 (0.7078), demonstrating better recognition of the minority emotion classes. The HybridT also performed competitively, reaching 89.57% accuracy and 0.6924 macro-F1 with substantially fewer parameters than the pretrained transformer backbones – outperforming the best conventional model by approximately 2.5 percentage points in accuracy and 0.05 in macro-F1 despite using no ImageNet pretraining. Consistent with the architectural rationale discussed in Section 5.1.2, these results confirm that self-attention-based feature extraction generalizes more effectively than convolutional or classical baselines on the fine-grained, highly imbalanced facial emotion recognition task posed by the ASD-FER dataset.

### 5.3. Per-Emotion Analysis

Table 11 presents the class-wise precision, recall, and F1-score of the five traditional models (AlexNet is omitted as it collapsed to a majority-class predictor), while Table 12 presents the class-wise precision, recall, and F1-score of the three transformer-based models on the held-out test set. Furthermore, Fig. 10 and Fig. 11 show the corresponding confusion matrices.

**Figure 10:**
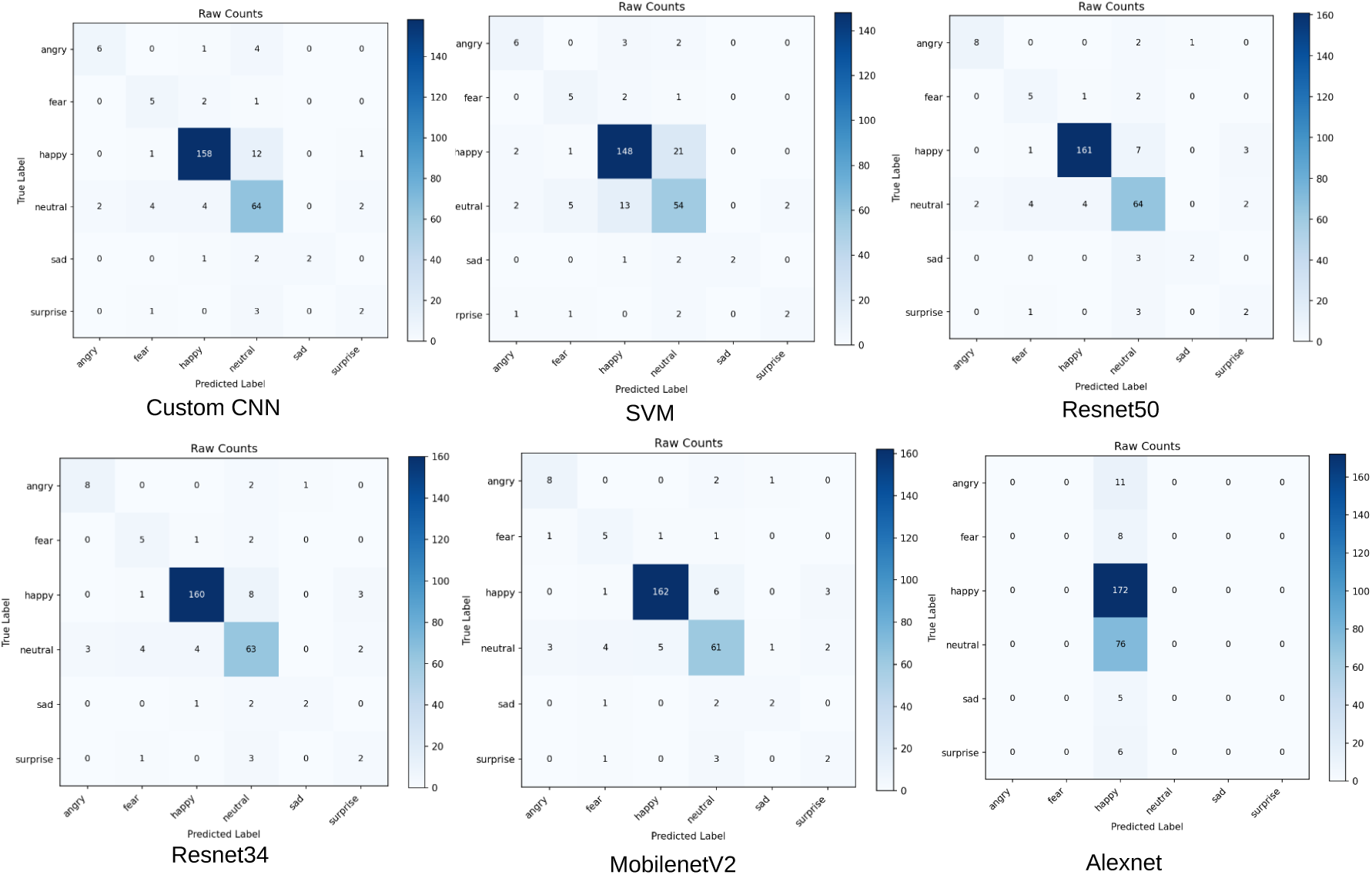
Confusion matrices for the six traditional models on the 278-image held-out test set.

**Figure 11:**
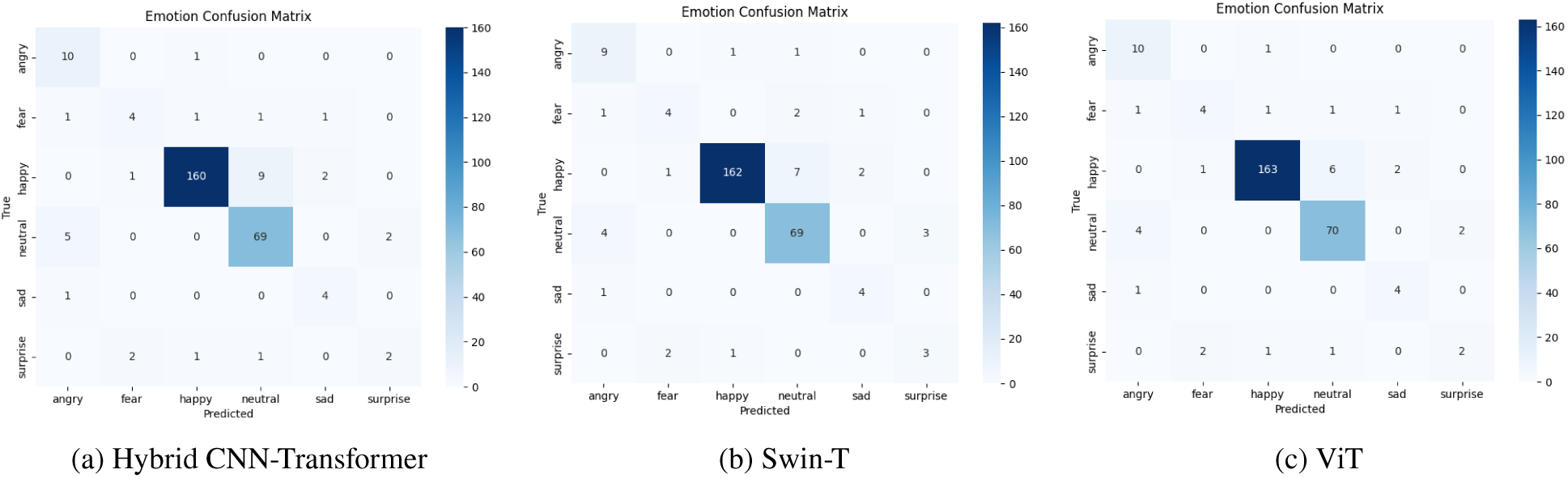
Confusion matrices for the three transformer-based models on the 278-image held-out test set. Each matrix shows true emotion labels (rows) against predicted labels (columns), with color intensity indicating sample count.

**Table 11.** Class-wise performance comparison of the five benchmark models on the held-out (90/10) ASD-FER test set. AlexNet is omitted as it collapsed to a majority-class predictor. Best value for each emotion and metric is shown in bold.

| Emotion | Custom CNN |  |  | SVM |  |  | ResNet34 |  |  | ResNet50 |  |  | MobileNetV2 |  |  | Support |
| --- | --- | --- | --- | --- | --- | --- | --- | --- | --- | --- | --- | --- | --- | --- | --- | --- |
|  | P | R | F1 | P | R | F1 | P | R | F1 | P | R | F1 | P | R | F1 |  |
| Angry | 0.7500 | 0.5455 | 0.6316 | 0.5455 | 0.5455 | 0.5455 | 0.7273 | **0.7273** | 0.7273 | **0.8000** | **0.7273** | **0.7619** | 0.6667 | **0.7273** | 0.6957 | 11 |
| Fear | **0.4545** | **0.6250** | **0.5263** | 0.4167 | **0.6250** | 0.5000 | **0.4545** | **0.6250** | **0.5263** | **0.4545** | **0.6250** | **0.5263** | 0.4167 | **0.6250** | 0.5000 | 8 |
| Happy | 0.9518 | 0.9186 | 0.9349 | 0.8862 | 0.8605 | 0.8732 | 0.9639 | 0.9302 | 0.9467 | **0.9699** | 0.9360 | 0.9527 | 0.9643 | **0.9419** | **0.9529** | 172 |
| Neutral | 0.7442 | **0.8421** | 0.7901 | 0.6585 | 0.7105 | 0.6835 | 0.7875 | 0.8289 | 0.8077 | 0.7901 | **0.8421** | **0.8153** | **0.8133** | 0.8026 | 0.8079 | 76 |
| Sad | **1.0000** | **0.4000** | **0.5714** | **1.0000** | **0.4000** | **0.5714** | 0.6667 | **0.4000** | 0.5000 | 0.6667 | **0.4000** | 0.5000 | 0.5000 | **0.4000** | 0.4444 | 5 |
| Surprise | 0.4000 | **0.3333** | 0.3636 | **0.5000** | **0.3333** | **0.4000** | 0.2857 | **0.3333** | 0.3077 | 0.2857 | **0.3333** | 0.3077 | 0.2857 | **0.3333** | 0.3077 | 6 |

**Table 12.** Class-wise performance comparison of the three transformer-based models on the held-out (90/10) ASD-FER test set. Best value for each emotion and metric is shown in bold.

| Emotion | HybridT |  |  | Swin-T |  |  | ViT |  |  | Support |
| --- | --- | --- | --- | --- | --- | --- | --- | --- | --- | --- |
|  | Precision | Recall | F1 | Precision | Recall | F1 | Precision | Recall | F1 |  |
| Angry | 0.6250 | **0.9091** | **0.7407** | 0.6000 | 0.8182 | 0.6923 | **0.6250** | **0.9091** | **0.7407** | 11 |
| Fear | **0.5714** | **0.5000** | **0.5333** | **0.5714** | **0.5000** | **0.5333** | **0.5714** | **0.5000** | **0.5333** | 8 |
| Happy | 0.9753 | 0.9186 | 0.9461 | **0.9878** | 0.9419 | 0.9643 | 0.9819 | **0.9477** | **0.9645** | 172 |
| Neutral | 0.8415 | 0.9079 | 0.8734 | 0.8734 | 0.9079 | 0.8903 | **0.8974** | **0.9211** | **0.9091** | 76 |
| Sad | **0.5714** | **0.8000** | **0.6667** | **0.5714** | **0.8000** | **0.6667** | **0.5714** | **0.8000** | **0.6667** | 5 |
| Surprise | **0.5000** | 0.3333 | 0.4000 | **0.5000** | 0.5000 | **0.5000** | **0.5000** | **0.3333** | 0.4000 | 6 |

Looking at the results of the five conventional models, it’s clear that how well each model does depends a lot on how many examples of each emotion it saw during training. For emotions with lots of samples, like Happy (172) and Neutral (76), all five models perform strongly, with high precision, recall, and F1 scores. For the rare emotions, the picture is different: with only five or six test images for Sad and Surprise, a single correct or incorrect prediction shifts the scores substantially, and recall in fact ties across all five models for Fear (0.6250), Sad (0.4000), and Surprise (0.3333) — every model recovers the same number of samples, and only precision separates them. One interesting finding is that Custom CNN, ResNet34, and ResNet50 all tie for the best Fear recognition (F1 = 0.5263), a tricky emotion since it can look similar to Angry or Neutral, while ResNet50 is strongest on Angry (F1 = 0.7619). MobileNetV2 comes out on top for Happy (F1 = 0.9529) and ResNet50 for Neutral (F1 = 0.8153), matching their strong overall performance seen earlier. Surprise is now the weakest class for every model, with recall stuck at 0.3333 despite its visually distinctive appearance, suggesting that six test samples are simply too few for reliable measurement rather than that the expression is inherently hard. Overall, the model we choose seems to matter most for the emotions that are hard to tell apart, like Fear versus Angry or Neutral, while performance on the rare emotions mostly comes down to not having enough data rather than which model was used. The confusion matrices support the quantitative results: all five models correctly identify most Happy and Neutral faces, but the rarer emotions are frequently absorbed into the Neutral column, consistent with Sad and Surprise showing the same recall (0.4000 and 0.3333) across every model. AlexNet’s matrix stands out the most, since every single image is predicted as Happy, showing that it failed to learn the other emotions at all. Among the remaining models, ResNet50 and MobileNetV2 show the cleanest diagonals overall, which lines up with their leading accuracy and balanced accuracy reported earlier, while the SVM shows the weakest separation between Happy and Neutral, consistent with its lower precision and recall on both classes (Happy F1 = 0.8732, Neutral F1 = 0.6835 — the lowest of the five models on both).

Furthermore, all three transformer models demonstrated similar trends across the six emotion classes. The *Happy* class was consistently recognized with the highest performance, achieving F1-scores above 0.94 for all models, which can be attributed to its large number of training samples (172 images) and its distinctive facial characteristics. The *Neutral* class also achieved strong performance, with ViT obtaining the highest F1-score (0.9091), followed closely by Swin-T (0.8903) and the HybridT (0.8734). These results indicate that the dominant classes in the dataset are learned reliably by all transformer architectures.

In contrast, the minority emotion classes remained more challenging. *Fear*, *Sad*, and *Surprise* achieved noticeably lower F1-scores across all models. Although the *Sad* class obtained a relatively high recall (0.80), its performance should be interpreted cautiously because only five test samples were available. Similarly, the *Fear* and *Surprise* classes contained only eight and six samples, respectively, making them more susceptible to prediction errors. The limited number of training examples, together with the subtle facial differences between these emotions, likely contributed to the reduced recognition performance.

Among the three models, ViT achieved the best overall performance for the *Happy* and *Neutral* classes, while Swin-T produced the highest F1-score for the *Surprise* class. The HybridT achieved competitive results despite having only 2.66M parameters, demonstrating that an efficient hybrid architecture can provide strong class-wise performance with substantially lower computational complexity than larger transformer backbones.

The confusion matrices in Fig. 11 further illustrate that most misclassifications occur among the minority emotion classes rather than between the dominant classes. Similar observations have been reported in general FER literature, where underrepresented emotions such as *Fear* and *Surprise* are typically more difficult to distinguish. In the ASD-FER dataset, this challenge is further amplified by the limited number of samples and the subtle, atypical facial expressions exhibited by children with ASD.

### 5.4. Comparison with Prior Work

Table 13 compares the proposed ASD-FER with representative ASD-specific and general facial emotion recognition (FER) studies, extending the qualitative comparison introduced in Section 2 with quantitative accuracy figures reported in each original study.

**Table 13.**
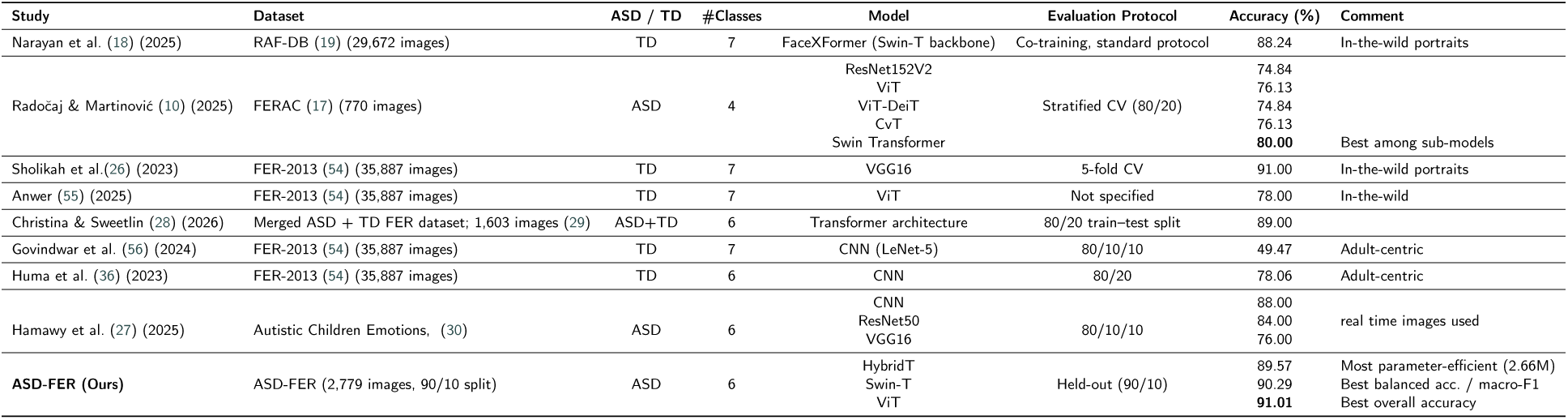
Comparison of the proposed ASD-FER with representative prior work on facial emotion recognition, spanning both ASD-specific and general (TD) studies. Results are reported as published in the original studies; differences in datasets, number of emotion classes, and evaluation protocols should be considered when interpreting cross-study comparisons.

Among the ASD-specific studies, the proposed approach achieves competitive performance across all three transformer-based configurations. The best-performing configuration, ViT, achieved 91.01% accuracy, while Swin-T obtained the highest balanced accuracy and macro-F1 score (0.7447 and 0.7078, respectively). The more compact HybridT reached 89.57% accuracy—trailing the two larger backbones by a modest margin while using only 2.66M parameters, roughly 17× and 39× fewer than Swin-T and ViT, respectively. These results compare favorably with previous ASD-specific studies, including the Swin Transformer reported by authors of (10) on the FERAC dataset (80.00% accuracy) and the ResNet50 model of Hamawy *et al.* (27) (88.00% accuracy). However, direct comparisons should be interpreted with caution because the studies differ in dataset size, number of emotion classes, and evaluation protocols. For example, FERAC contains 770 images with four emotion classes, whereas the proposed ASD-FER dataset contains six emotion classes and is evaluated using an independent held-out (90/10) protocol.

Compared with general-population FER studies, the proposed method also demonstrates competitive performance. Recent transformer-based methods such as FaceXFormer (18) achieved 88.24% accuracy on RAF-DB, while VGG16-based methods reported approximately 91.00% accuracy on FER-2013 under five-fold cross-validation. Although these studies report similar or slightly higher accuracies, they are evaluated on substantially larger, adult-centric datasets with different characteristics from ASD facial expression datasets. Therefore, rather than making direct state-of-the-art claims across different benchmarks, the results indicate that ASD-FER provides strong performance for facial emotion recognition in children with ASD, achieves results that exceed comparable ASD-specific prior work, and remains competitive with methods developed for general-population FER tasks.

## 6. Grad-CAM Interpretability Analysis

### 6.1. Grad-CAM Methodology

Gradient-weighted Class Activation Mapping (Grad-CAM) was employed to visualize the image regions that contributed most to the predicted emotion for all three transformer-based models. Grad-CAM computes the gradients of the predicted class score *y^c^* with respect to the feature maps *A^k^* of a selected target layer, producing a weighted activation map:

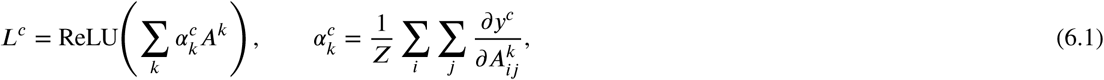

where *α^c^* denotes the global average gradient weight for class *c*, and *Z* is the number of spatial locations. The ReLU operation retains only the positive contributions that support the predicted class.

Different target layers were selected according to each model architecture. For the **HybridT**, Grad-CAM was applied to the final BatchNorm2d layer of the CNN stem (cnn.layers[3]), which preserves a spatial feature map of size [*B*, 128, 56, 56] before transformer encoding. For the **Swin Transformer**, the LayerNorm layer of the final Swin block (backbone.layers[-1].blocks[-1].norm1) was used, with the resulting features reshaped from [*B*, *H*, *W*, *C*] to [*B*, *C*, *H*, *W*]. For the **ViT** model, which does not produce convolutional feature maps, the LayerNorm preceding the final transformer block (backbone.encoder.layer[-1].layernorm_before) served as the target layer. The output token sequence [*B*, 197, 768] was converted into a 14 × 14 spatial feature map by discarding the class token and reshaping the remaining patch embeddings to [*B*, 768, 14, 14].

The generated heatmaps follow a consistent color convention: **red** and **yellow** indicate regions that contribute most strongly to the predicted emotion, **green** represents moderate importance, and **blue** denotes minimal contribution. The heatmaps were qualitatively examined to verify whether the models focused on psychologically meaningful facial regions, particularly the eyes, eyebrows, and mouth, the primary regions humans use to decode facial expressions (5; 6).

### 6.2. Visualization Results per Model

#### 6.2.1. Hybrid CNN-Transformer

Fig. 12 shows Grad-CAM outputs for the HybridT across all six emotion classes. The heatmaps exhibit relatively low and diffuse activation magnitudes compared to the pretrained backbones, consistent with the model being trained from scratch on a much smaller corpus. Despite this, the heatmaps consistently highlight the eyes, eyebrows, and mouth across all classes (e.g., *angry*, *sad*, *fear*), indicating that even with limited activation strength, the model’s discriminative evidence is anchored in psychologically meaningful facial areas rather than background or irrelevant regions.

**Figure 12:**
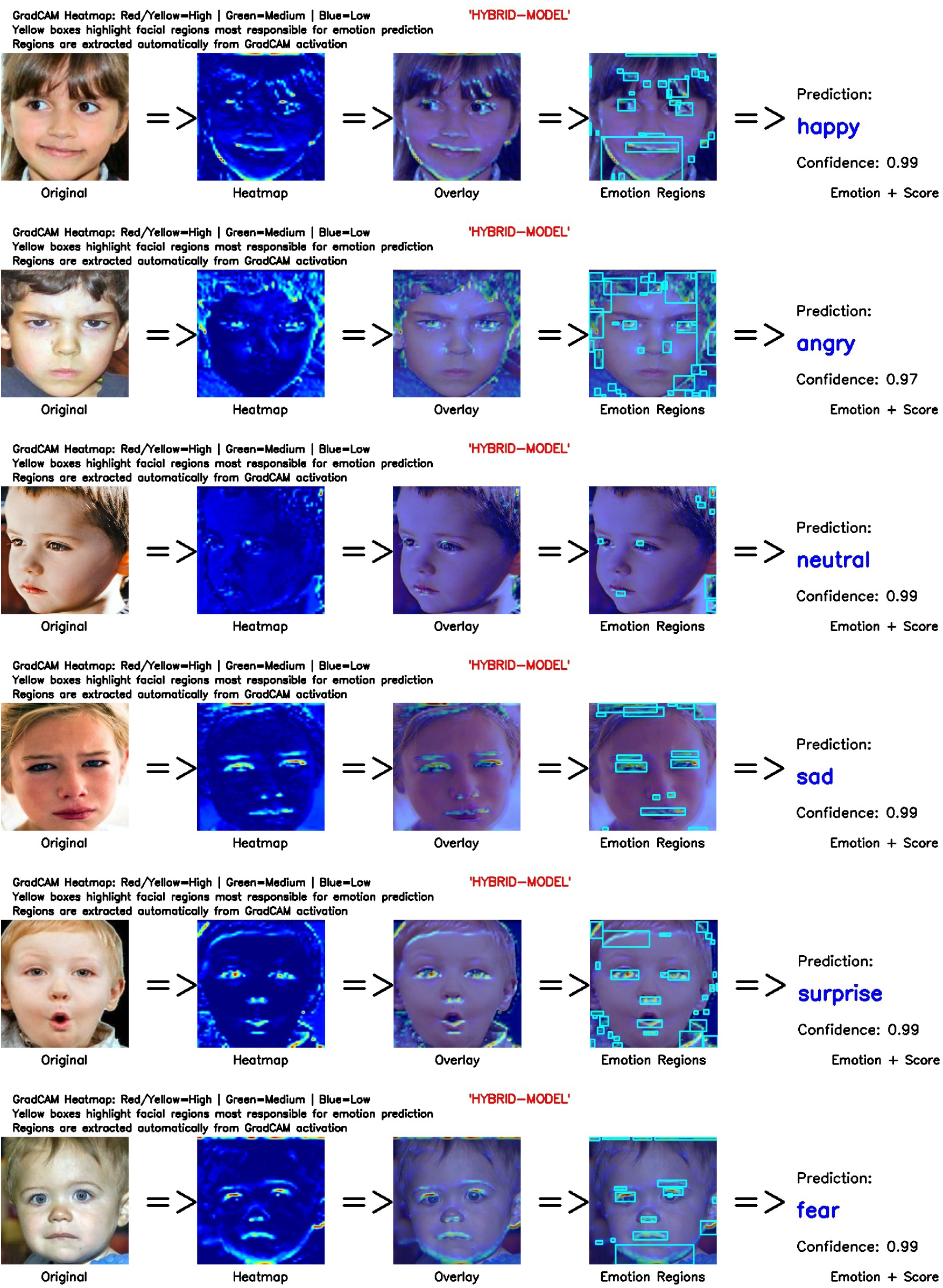
Grad-CAM visualizations generated by the HybridT for representative facial emotion samples. Red and yellow regions indicate the most discriminative facial areas contributing to the predicted emotion. The facial images are from the publicly available Kaggle dataset, Facial_data_ASD, by Md. Shafiul Alam (34).

#### 6.2.2. Swin-T

Fig. 13 presents Grad-CAM visualizations for the Swin Transformer. Activation is markedly stronger and more spatially concentrated than the HybridT model, with red/yellow hotspots tightly clustered around the central face, particularly the eyes, eyebrows, and mouth (e.g., *surprise*, *fear*, *angry*). This tight, high-contrast localization supports the hierarchical window-based attention of Swin-T, which appears to consistently suppress peripheral and background regions in favor of the core expressive facial features.

**Figure 13:**
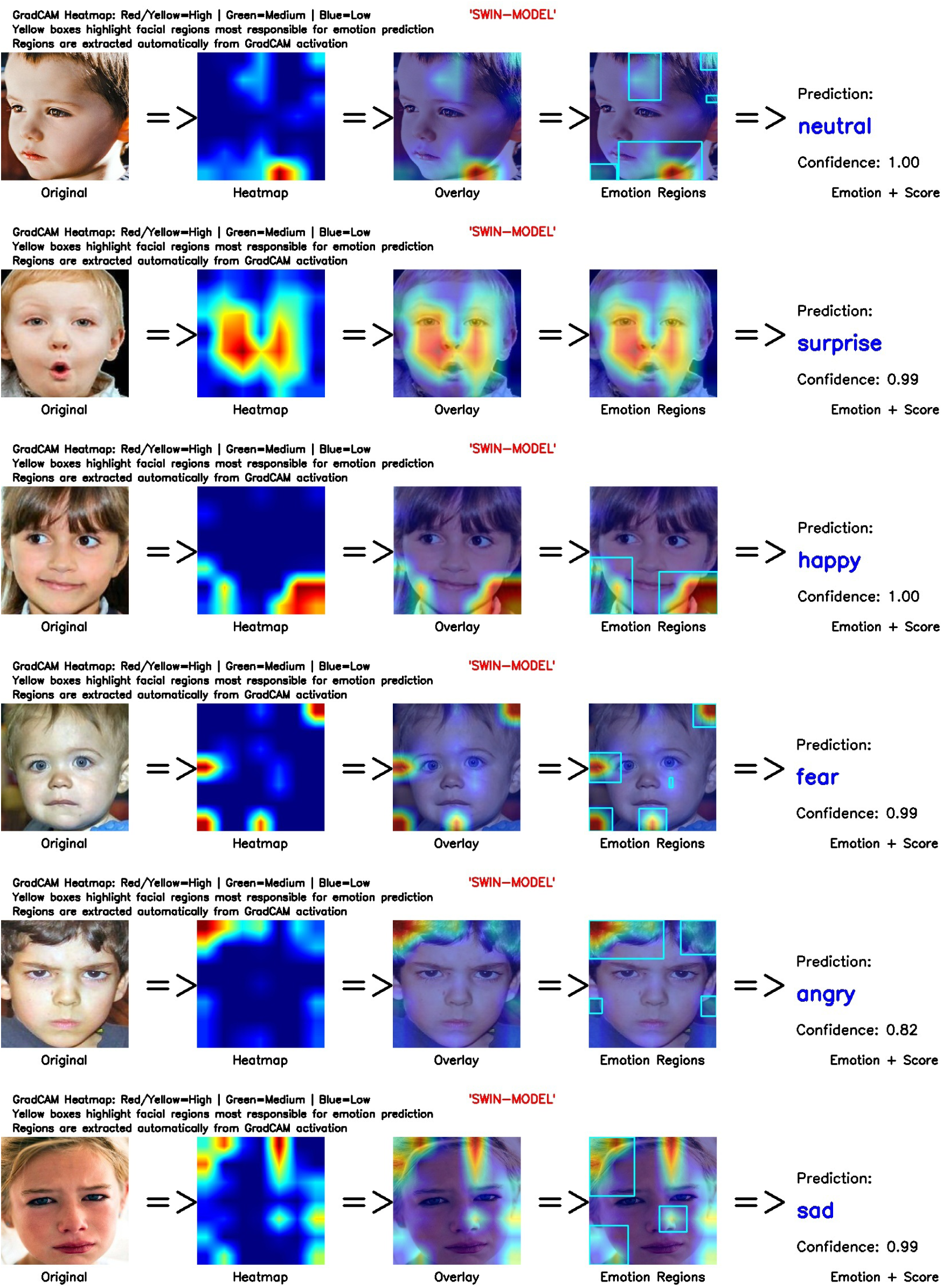
Grad-CAM visualizations generated by the Swin-T for representative facial emotion samples. Red and yellow regions indicate the most discriminative facial areas contributing to the predicted emotion. The facial images are derived from the publicly available Kaggle dataset, Facial_data_ASD, by Md. Shafiul Alam (34).

#### 6.2.3. Vision Transformer (ViT)

Fig. 14 shows the corresponding results for ViT. While activation still centers on the eyes, eyebrows, and mouth, the heatmaps are visibly broader and more distributed across the face than Swin-T, with secondary activation extending to the forehead and cheek regions (e.g., *angry*, *sad*). This is consistent with ViT’s global self-attention mechanism, which integrates evidence across the entire patch sequence rather than emphasizing a compact local neighborhood, at the cost of somewhat less spatially precise localization than Swin-T.

**Figure 14:**
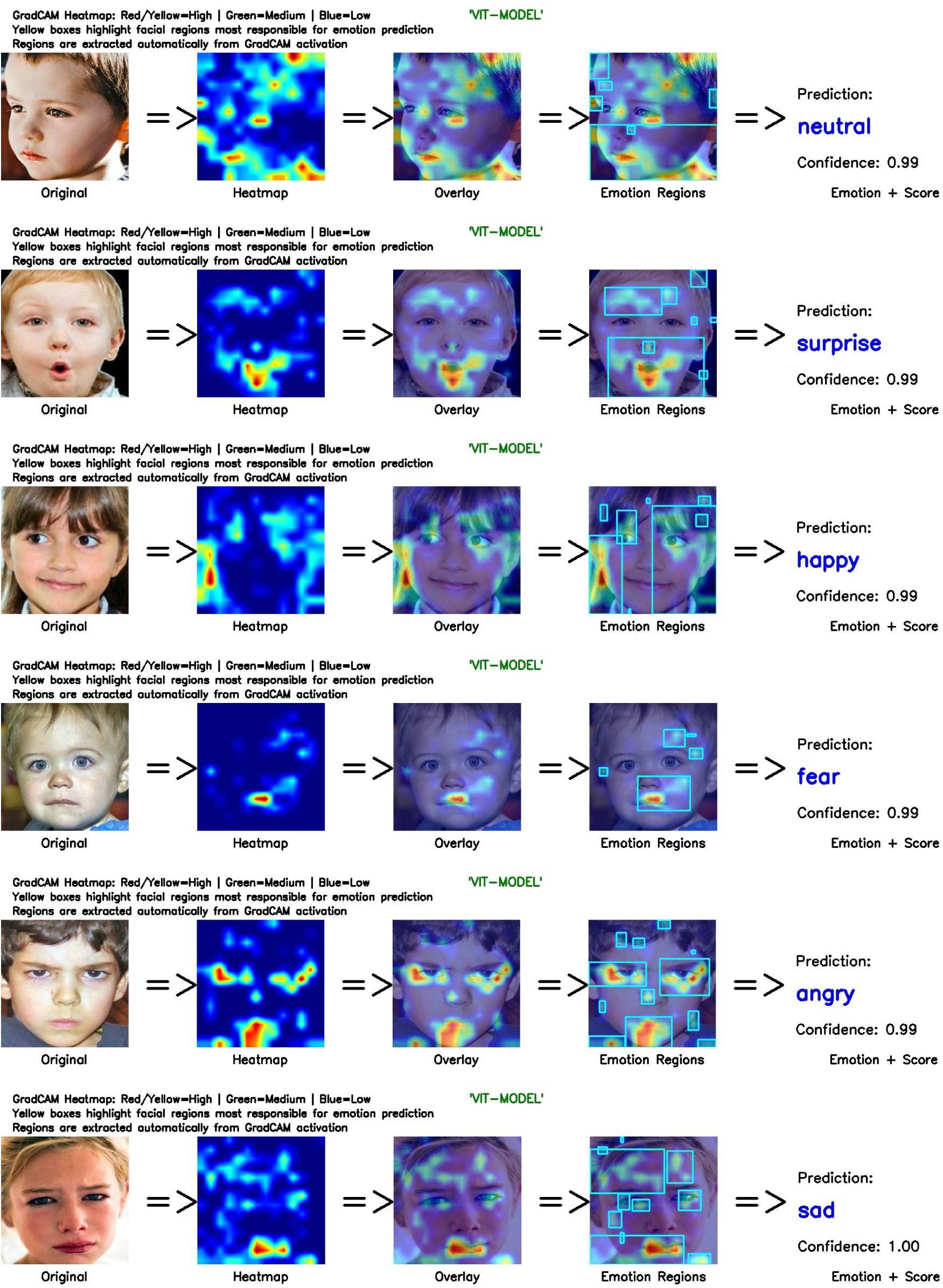
Grad-CAM visualizations generated by the ViT for representative facial emotion samples. Red and yellow regions indicate the most discriminative facial areas contributing to the predicted emotion. The facial images are derived from the publicly available Kaggle dataset, Facial_data_ASD, by Md. Shafiul Alam (34).

**Figure 15:**
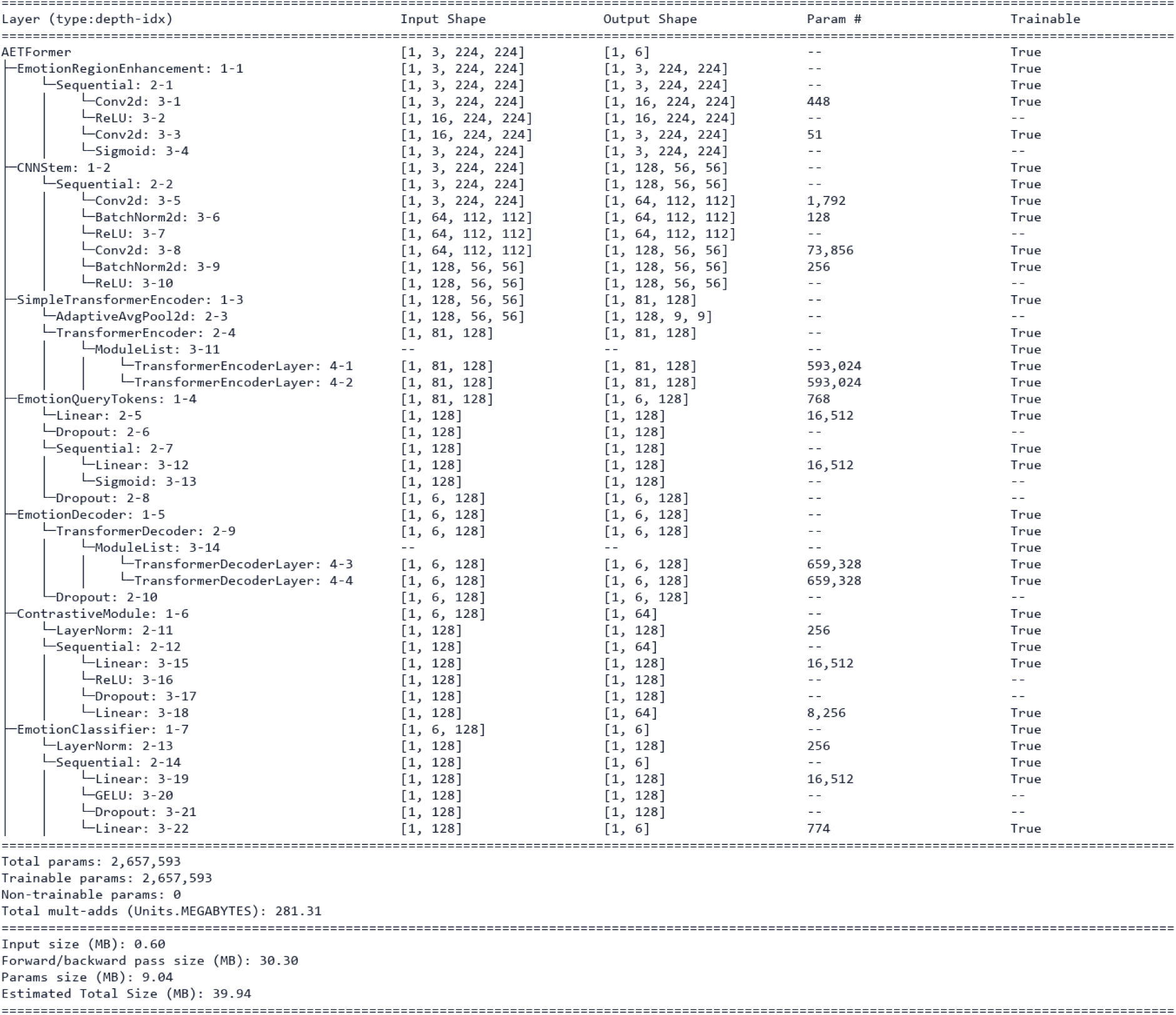
Layer-wise architecture summary of the HybridT.

**Figure 16:**
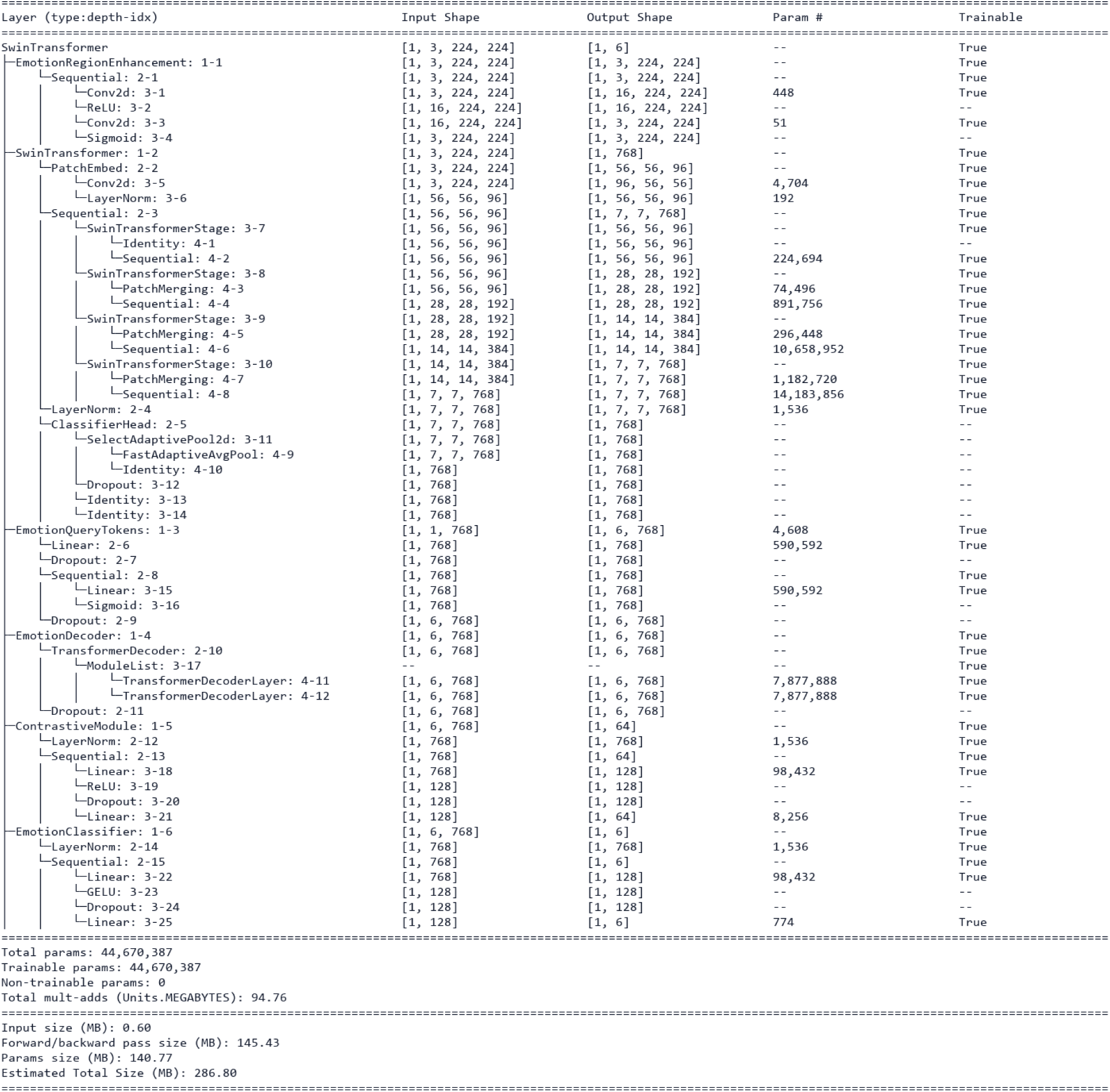
Layer-wise architecture summary of the Swin Transformer (Swin-T).

**Figure 17:**
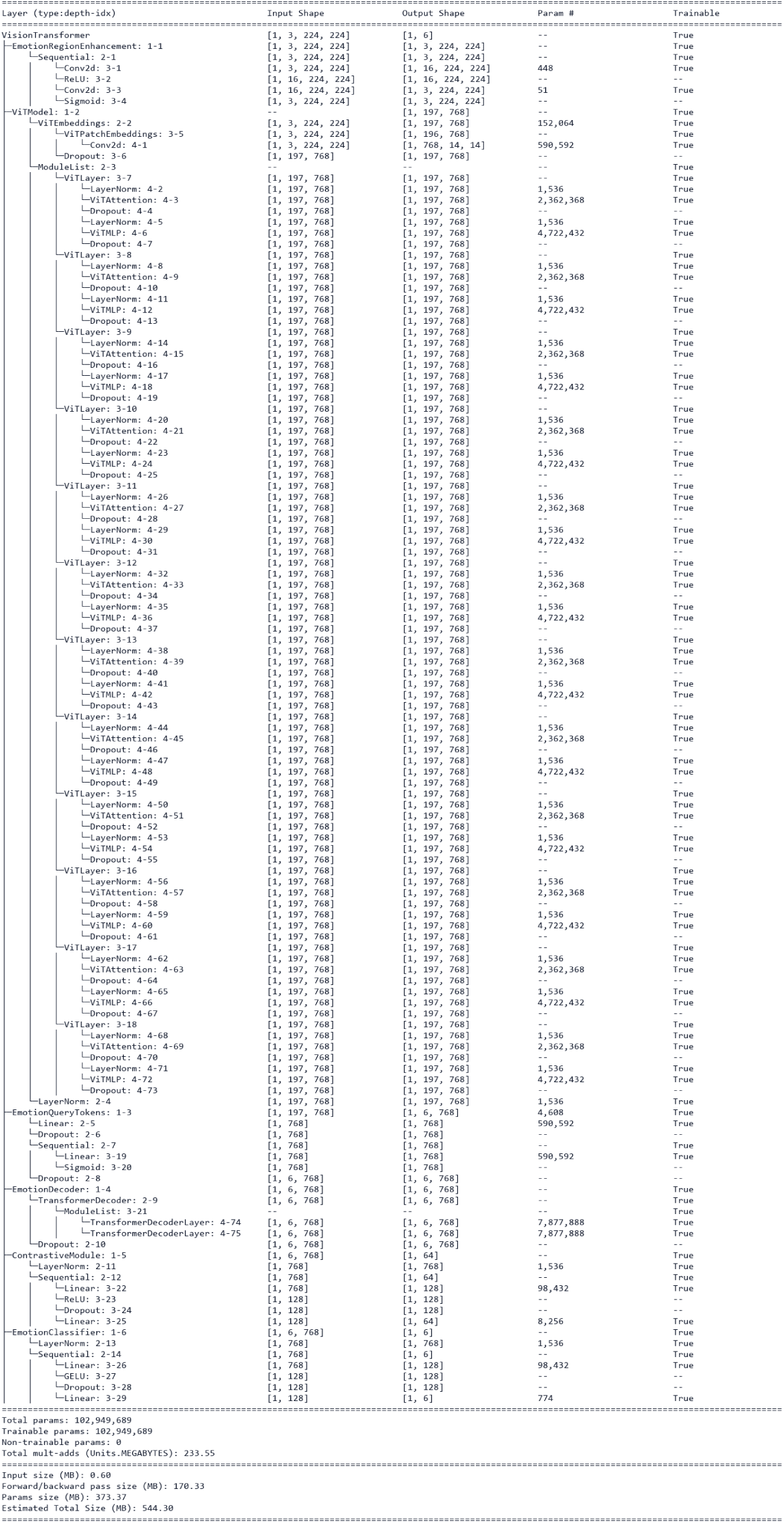
Layer-wise architecture summary of the ViT.

## 7. Limitations and Future Work

Although ASD-FER and the accompanying models show strong results, several limitations should be noted.

The dataset contains 2,779 images, which remains small relative to large-scale general FER benchmarks, and exhibits class imbalance across fear, sadness, and surprise categories. Images were sourced from publicly available online material rather than clinical recruitment. Future datasets built through clinical collaborations — with confirmed diagnoses, longitudinal observations, and richer subject metadata — would substantially improve reliability and generalizability.

This study focuses on static images, whereas emotions naturally unfold over time. Extending to video sequences and multimodal cues such as gaze, speech, and physiological signals is a natural next step. The fixed six-emotion taxonomy ensures consistency with established FER benchmarks but may not fully capture the diversity and atypicality of emotional expressions in children with ASD.

Regarding annotation, disagreements were resolved through direct discussion among the three raters rather than by an independent adjudicator, which may have introduced anchoring bias in borderline cases. The inter-rater statistics in Section 3.2.4 were computed on pre-discussion labels and are unaffected by this. Future annotation efforts should consider a fully blinded resolution step, such as an independent fourth rater adjudicating without knowledge of the other raters’ responses.

Because subject identities in the source images are unverified, fold splitting was performed at the original-image level rather than a confirmed subject (child) level. If any child appears across multiple distinct source photographs, strict subject-level independence between folds cannot be guaranteed, even though augmentation-derived leakage within a fold is already prevented. Clinically recruited data with verified subject identifiers would allow rigorous subject-level splitting and eliminate this uncertainty.

Finally, Grad-CAM provides useful visual explanations but remains a post-hoc technique. Complementing it with additional explainability methods and clinical expert validation is an important direction for future work.

## 8. Conclusion

This paper presented ASD-FER, an annotated multi-class benchmark for facial emotion recognition in children with Autism Spectrum Disorder, together with a unified comparative evaluation of nine machine learning and deep learning models. Three complementary transformer paradigms — HybridT, Swin-T, and ViT — were evaluated under an identical preprocessing, training, and evaluation framework to enable a controlled architectural comparison. We refer to this shared query-based decoding architecture as **EQFormer**, evaluated here with three backbone configurations (HybridT, Swin-T, ViT) on ASD-FER. As EQFormer has so far been validated only on this dataset, an extended version of this work will assess its generalization to general-population FER benchmarks and additional ASD-specific datasets. Results from stratified 10-fold cross-validation and an independent held-out evaluation showed that the transformer-based configurations generally outperformed the conventional CNN-based approaches under the evaluated protocol. Among the evaluated models, ViT achieved the highest overall accuracy, while Swin-T produced the best balanced accuracy and macro-F1 score, indicating stronger recognition of minority emotion classes. Grad-CAM visualizations further showed that all three transformer models focused primarily on the eyes, eyebrows, and mouth rather than background or irrelevant regions, providing qualitative evidence that the models attend primarily to facially relevant regions. Overall, the proposed benchmark and evaluation provide a foundation for future ASD-specific facial emotion recognition research and may inform the development of more accurate, interpretable, and practically deployable assistive systems. The ASD-FER dataset and full experimental source code are publicly available at https://github.com/HAIM-LAB/ASD-FER.

## **A.** Fold-by-Fold Cross-Validation Results

Table 14 presents the complete per-fold accuracy, balanced accuracy, precision, recall, weighted F1, and macro-F1 scores for the three transformer-based models across all 10 folds of the stratified cross-validation protocol.

**Table 14:**
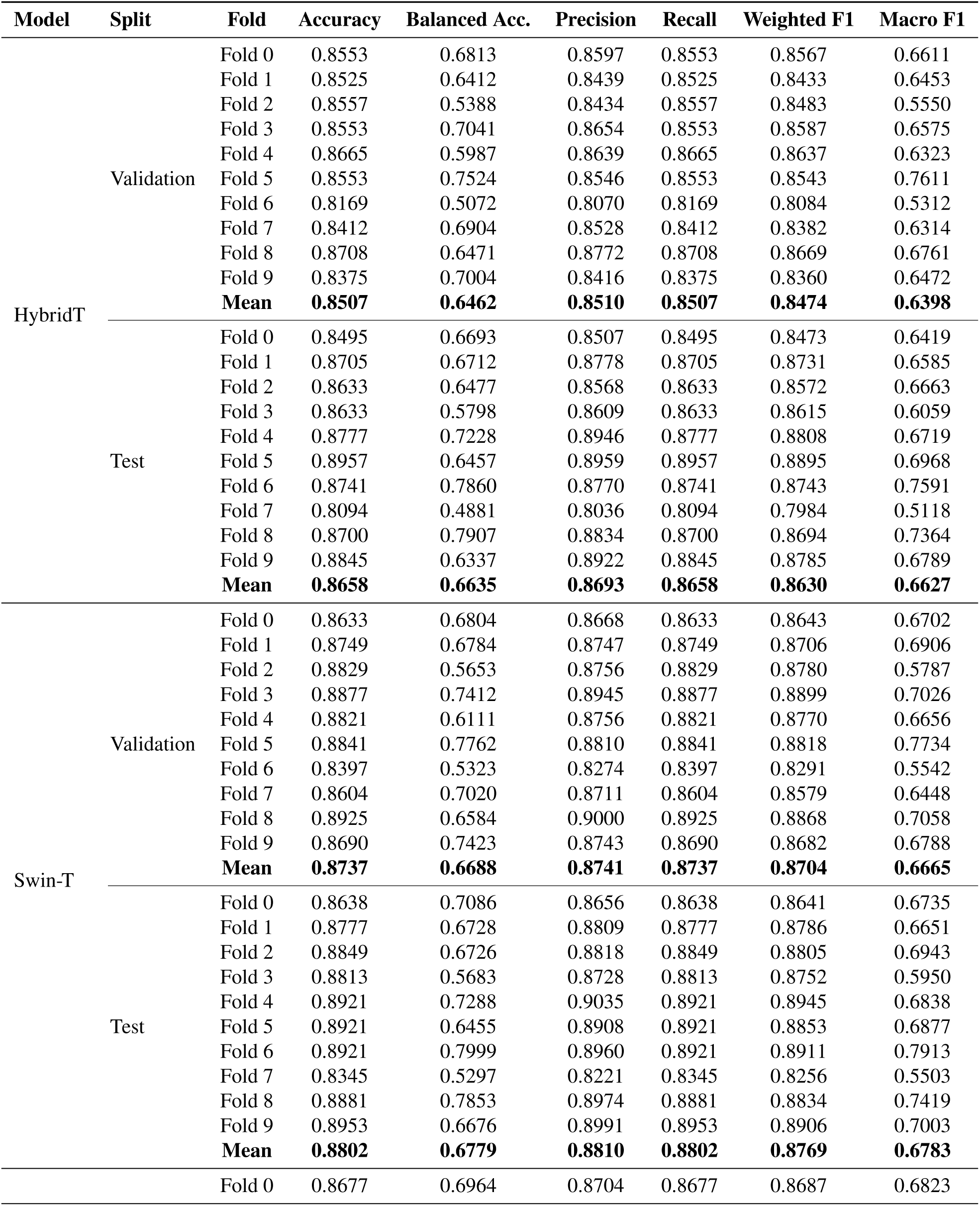
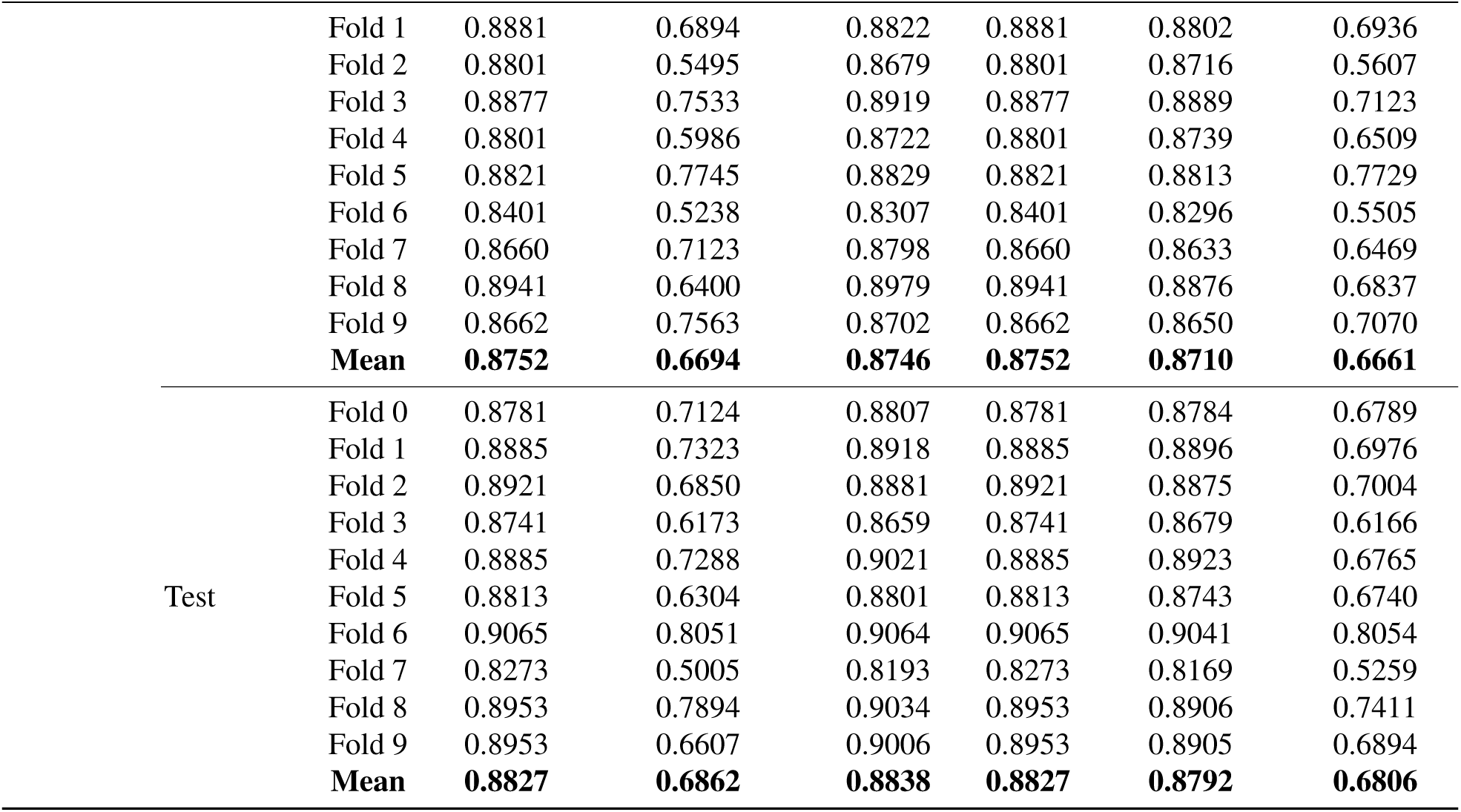
Fold-by-fold stratified 10-fold cross-validation results for the Hybrid CNN-Transformer, Swin-T, and ViT models on validation and test splits.

## **B.** Model Architecture Summaries

This appendix provides the complete layer-wise architecture summaries for the three transformer-based models evaluated in this study, generated using torchinfo, including per-layer output shapes, parameter counts, and trainable status.

## Acknowledgment

This research is supported in part by the Public University Partnership Program at the Louisiana Department of Health, Bureau of Health Services Financing, through the National Institutes of Health (NIH) Award No. 10000411, Institutional Development Award (IDeA) from the National Institute of General Medical Sciences of the National Institutes of Health under grant number P20 GM103424-21, and the Data Science Block Grant from the National Institute of General Medical Sciences, NIH Grant No. 2P20GM103424-24.

## Declaration of Generative AI and AI-Assisted Technologies in the Manuscript Preparation Process

During the preparation of this work, the authors used Claude and ChatGPT for assistance with coding-related errors and debugging. Claude and Grammarly were used for text modification, grammar correction, and sentence restructuring. The authors reviewed and edited all AI-assisted outputs as needed and take full responsibility for the content of the published article.

## CRediT authorship contribution statement

**Shahana Yasmin Chowdhury:** Conceptualization, Methodology, Software, Validation, Data curation, Investigation, Formal analysis, Writing – original draft, Writing – review & editing. **Bithi Banik:** Methodology, Software, Validation, Investigation, Writing – original draft. **Tracey Ann Knaus:** Data curation, Writing – review & editing. **Christopher David Moore:** Data curation, Writing. **Henry Fordjour Ansah:** Data curation. **Huda Hammad:** Data curation. **Pranish Ghimire:** Data curation, review. **Md Tamjidul Hoque:** Methodology, Supervision, Writing – review & editing. **Shreya Banerjee:** Conceptualization, Supervision, Formal analysis, Writing – review & editing, Project administration, Funding acquisition.

